# A Canonical Neural Mechanism for Behavioral Variability

**DOI:** 10.1101/086074

**Authors:** Ran Darshan, William E. Wood, Susan Peters, Arthur Leblois, David Hansel

## Abstract

The ability to generate variable movements is essential for learning and adjusting complex behaviors. This variability has been linked to the temporal irregularity of neuronal activity in the central nervous system. However, how neuronal irregularity actually translates into behavioral variability is unclear. Here we combine modeling, electrophysiological and behavioral studies to address this issue. We demonstrate that a model circuit comprising topographically organized and strongly recurrent neural networks can autonomously generate irregular motor behaviors. Simultaneous recordings of neurons in singing finches reveal that neural correlations increase across the circuit driving song variability, in agreement with the model predictions. Analyzing behavioral data, we find remarkable similarities in the babbling statistics of 5-6 month-old human infants and juveniles from three songbird species, and show that our model naturally accounts for these ‘universal’ statistics.

## INTRODUCTION

Behavioral variability is a pivotal component of motor learning and adaptation^1-2^. While young individuals can usually produce non-stereotyped disorganized behaviors, motor exploration is more often expressed as movement variability around a stereotyped motor pattern. Highly irregular patterns of activity, which are ubiquitous in the brain^3^, are thought to underlie variable motor behaviors^4-5^. Specifically in songbirds, a neural circuit necessary for song learning in juveniles^6-7^ has been recently shown to be responsible for vocal variability both in adults^8^ and throughout development^7,9-10^. This circuit includes two cortical-like areas: a premotor nucleus, the lateral magnocellular nucleus of the anterior nidopallium (LMAN), and its efferent motor nucleus, the robust nucleus of the arcopalium (RA). While RA is essential in driving the effectors (muscles or muscle synergies) producing the song^11^, LMAN is not necessary for song production in adults^6-7^, but has a key role throughout development and in adults in driving variability in the song^9-10^and in the activity of RA neurons^12^.

The idea that temporal irregular activity of neurons in the central nervous system (CNS) is capable of generating behavioral variability may seem obvious. A careful examination, however, reveals that the link between irregularity in neural activity and behavioral variability is far from being straightforward. This is because to impact the behavior, patterns of activity generated in the central nervous system must also be *spatially* correlated (i.e., correlated across neurons). For example, consider the minimal model of a cortical network driving motor behavior depicted in Fig.1a. It consists of many neurons randomly connected recurrently, divided into D functional groups; each group is composed of M neurons (larger than D by over an order of magnitude) that project to one effector of the motor behavior. The collective dynamics of the network give rise to highly irregular firing patterns as a consequence of the interplay between excitation and inhibition^13^ (Fig.1b, Supplementary Fig.1a-b). Despite this large variability in their activity, unless the number of neurons in a group is very small, fluctuations in the effectors are negligible (the coefficient of variation of the input to the effector, CV_eff_, see Materials and Methods, is very small; Fig.1a, Supplementary Fig.1e). This stems from the fact that the network activity is only weakly correlated across neurons (on the order of 1/N, where N is the number of neurons in the network, Fig.1b right, Supplementary Fig.1d) and thus, by virtue of the law of large numbers, the fluctuations they induce in the net input to an effector “average out”. This example emphasizes the fact that for the fluctuations to be transferred robustly from the CNS to the effectors, neuronal firing in the motor network must be sufficiently correlated within a neural population projecting to the same effector.

**Figure 1.**
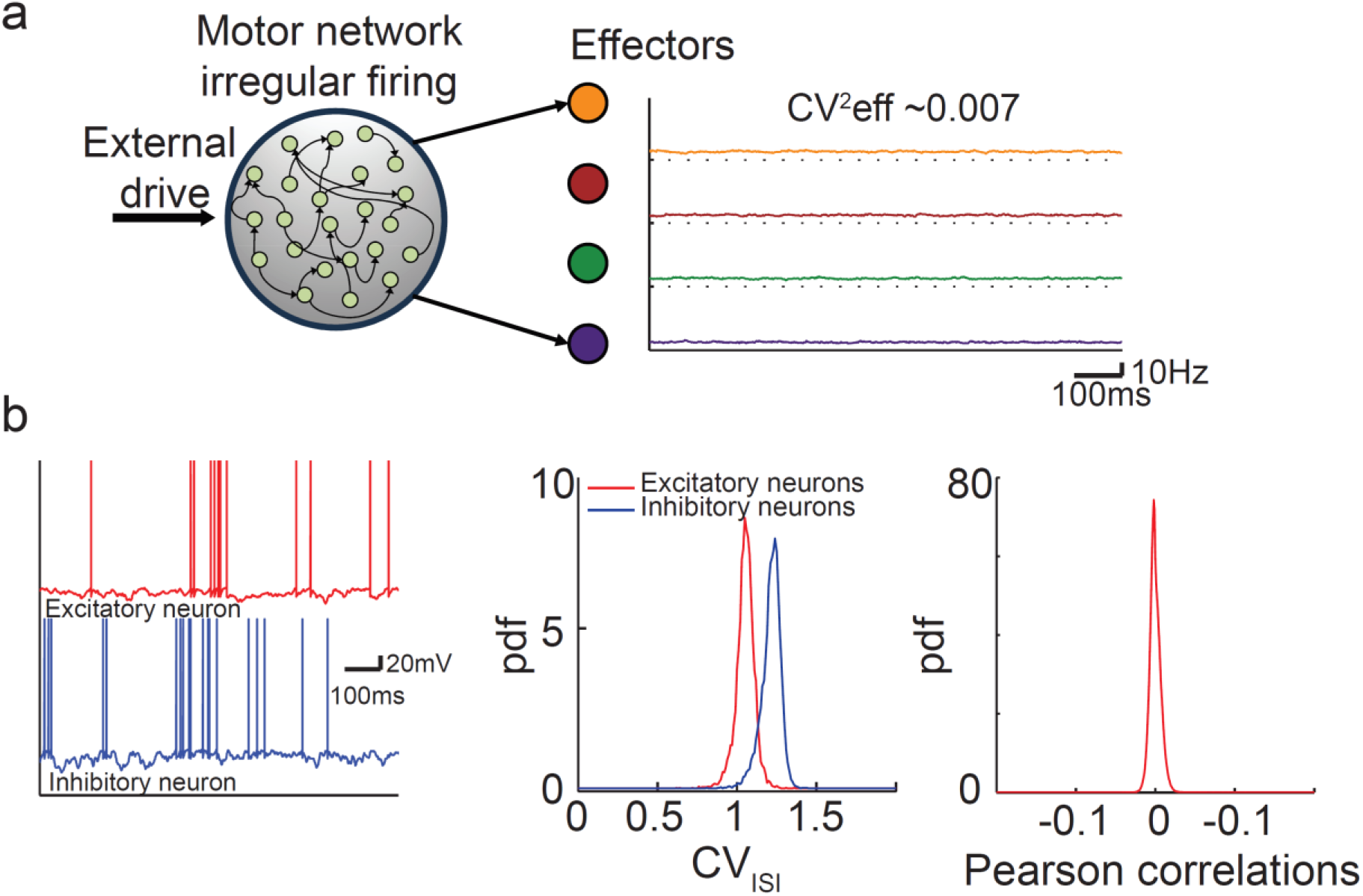
**Fluctuations in the inputs to the effectors are very weak when noise is generated autonomously in the motor network.a.** The motor network projects in a topographic manner to D effectors (D=10 effectors, 4 represented): each effector receives inputs from a different group of M=1000 neurons. In spite of the large variability of the neuronal activity, the variability of the effectors (right) is extremely small (coefficient of variation of the effector averaged over the 10 effectors: 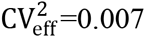). **b**. The neuronal activity in the motor network is highly irregularandthe correlations across neurons are tightly distributed around zero. Left: Voltage traces for one excitatory (E, red) and one inhibitory (I, blue) neuron. Middle: Distributions of coefficient of variation of the Inter-Spike-Interval, CV_ISI_. Right: Probability density function (pdf) of Pearson correlation coefficients in the network.

While the mechanism underlying *asynchronous* irregular spiking activity in recurrent networks of excitatory and inhibitory neurons is well understood^13-16^, how the CNS autonomously generates patterns of activity, which are both temporally irregular *and* correlated across neurons, remains an open fundamental question^16-19^. A key result of our theoretical work is that the activity of neurons in the motor network driving the effectors will be highly irregular and also spatially correlated if this network receives topographically organized excitatory projections from another upstream strongly recurrent network, hereafter premotor network. In the context of the circuit driving song variability in songbirds, our theory predicts that correlations emerge along the LMAN-RA circuit, namely that correlations across neurons are very weak in LMAN but substantial in RA. We validate this prediction with simultaneous extracellular recordings of neurons in singing finches. Our theory also suggests that vocal variability in different species of juvenile vocal learners should exhibit very similar statistics as a consequence of universal statistical properties of the circuit dynamics. We verify this prediction by comparing the statistics of the song produced during the babbling phase of three species of songbirds as well as of human infants. This work appeared in an abstract form^49^.

## RESULTS

### Motor variability emerges from the interplay between recurrent connections and topographic feedforward organization

We first show that temporally irregular and spatially correlated patterns of spiking activity can robustly emerge in a circuit of topographically organized and strongly recurrent networks. To this end, we consider the circuit depicted in Fig.2a. Neurons in the motor network which project to the same effector share a fraction, f, of their premotor inputs, and this shared component is different from one group to the other (see Materials and Methods for a detailed description of the architecture). With this architecture the spiking activities of the neurons in the premotor, as well as in the motor network, are highly irregular, as a result of their recurrent dynamics. There is, however, an important difference between the networks in the spatial structure of their activities. In the *premotor* network correlations across neurons are typically extremely weak (Fig.2b-c, Fig.4a). In contrast, in the *motor* network pairs of neurons projecting to the same effector are substantially and positively correlated, whereas correlations are weak (and possibly negative) for neurons projecting to different effectors (Fig.2d-g, Fig.4b and Supplementary Fig.3-5). These functional correlations are highly robust and only weakly influenced by the model parameters (Supplementary Fig.3d-e, Supplementary Fig.5d-f and Supplementary Information). As a result, fluctuations are amplified along the circuit (Fig.2e) and the variability is robustly transferred to the effectors.

**Figure 2.**
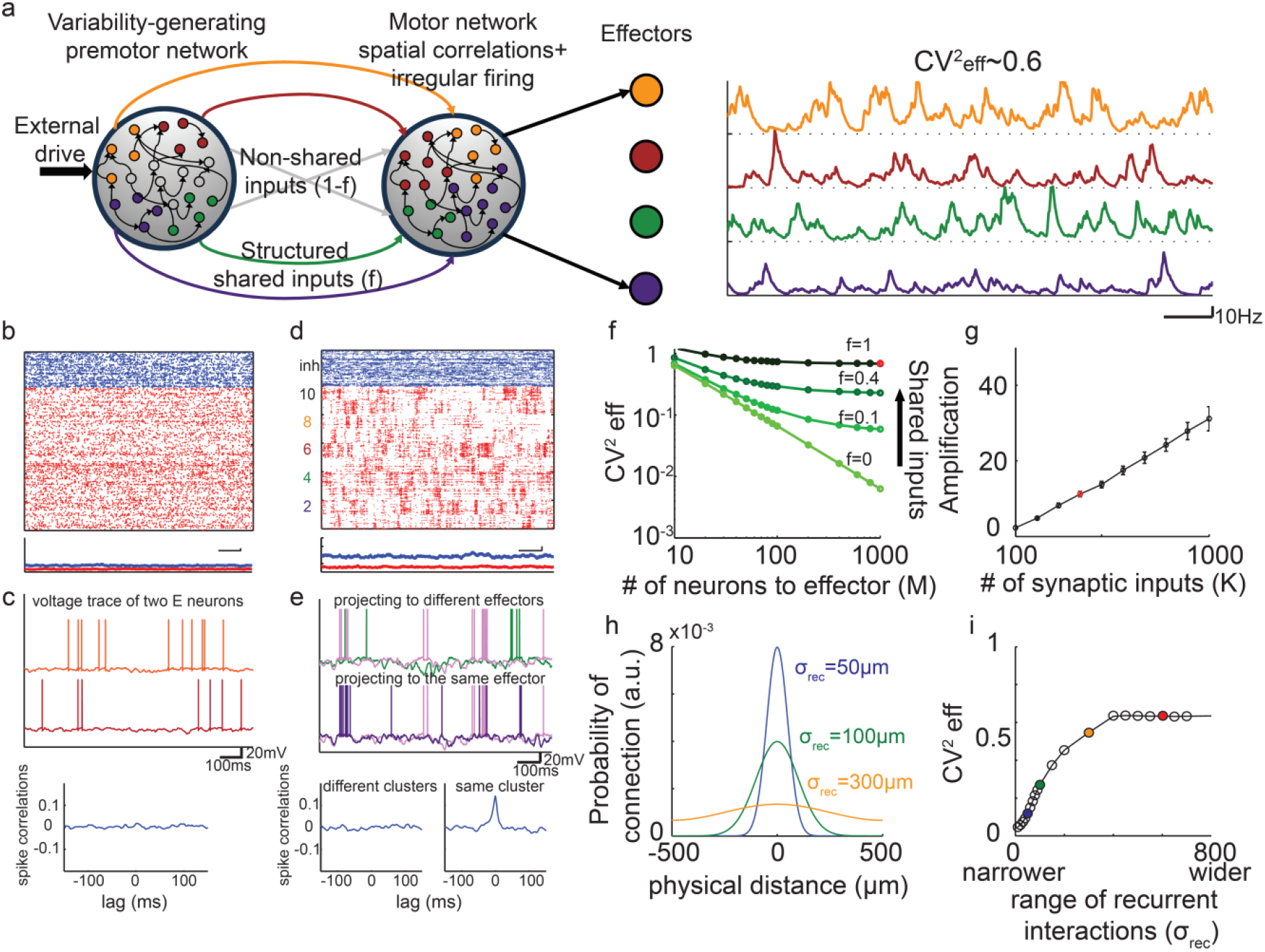
**Architecture and neuronal dynamics of a generic neural circuit driving behavioral variability. a.** When the premotor-to-motor projections are topographically organized, fluctuations in the inputs to the effectors are large. Left: the circuit architecture. In the motor network, neurons in the same group (same color, projecting to the same effector) share a fraction f of their premotor inputs (arrows colored identically to the corresponding group) and have a fraction 1-f of non-shared inputs (grey arrows). Right: the inputs to the effectors are highly variable 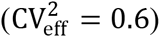. **b-c.** In the *premotor* network single neuron activity is highly irregular and very weakly correlated. **b**.Top: Raster plots of E (red) and I (blue) premotor neurons. Bottom: instantaneous mean activity of the E and I neurons (scale bar: 100ms and 10Hz). **c.** Voltage traces of two excitatory premotor neurons (top) and their spike crosscorrelations (bottom). **d-e.** In the *motor* network, single neuron activity is highly irregular *and* neurons are correlated. **d.** Top: Raster plots of E and I populations. Bottom: instantaneous mean activity of the E and I neurons (scale bar: 100ms and 10Hz).**e.** Voltage traces of two neurons in the motor network projecting to different (top) and same (bottom) effectors. Bottom: Pairs of neurons projecting to the same effector are substantially correlated (right); Pairs projecting to different effectors are very weakly correlated (left; see also Fig.4). **f.** The variability of the inputs to the effectors increases with the fraction of shared inputs and is substantial even if the number of inputs per effector, M, is large. This is because in the motor network the activities of the neurons belonging to the same group are correlated. **g.** The circuit amplifies fluctuations. The amplification factor, 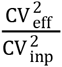, (see Materials and Methods) measures the ratio between the variability of the effectors 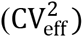 and of the input to the motor network 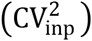. It increases linearly with the average number of synapses per neuron, K (mean±s.e.m; see also Supplementary Fig.3f). **h.** The connection probability of two neurons in the motor network depends on their distance (see Material and Methods) with a footprint *σ*_*rec*_. The diameter of the motor network is *λ* = 1000µm. 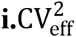 decreases when narrowing the footprint of the recurrent interactions in the motor network. Red dot in the figure corresponds to the parameters used in (a-e).

**Figure 3.**
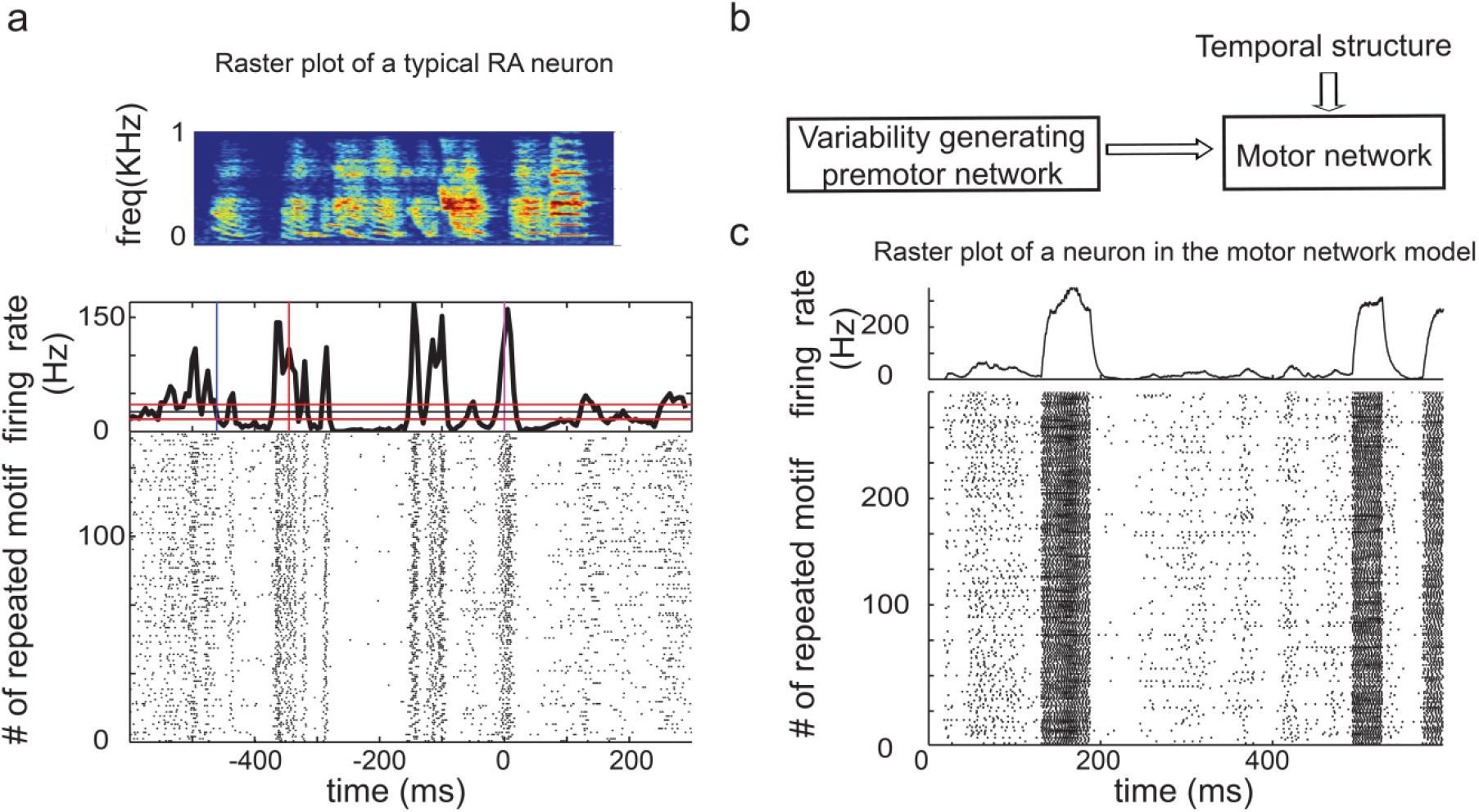
**Examples of single unit recordings in zebra finch RA nucleus and in the model motor network. a.** Top: song motif of a zebra finch. Bottom: recordings of RA single unit over 133 repetitions of song motif, aligned to one syllable in the motif (lower panel) and the corresponding average firing rate (upper panel; 5ms bin size). **b.** Extension of the model depicted in Fig.2a. Neurons in the motor network receives also *temporally structured* feedforward inputs, representing HVC inputs in the adult zebra finch (see main text and Material and Methods**)c.** Raster plot and corresponding average firing rate of a neuron in the motor network of the model circuit in the presence of temporally structured feedforward input.

**Figure 4.**
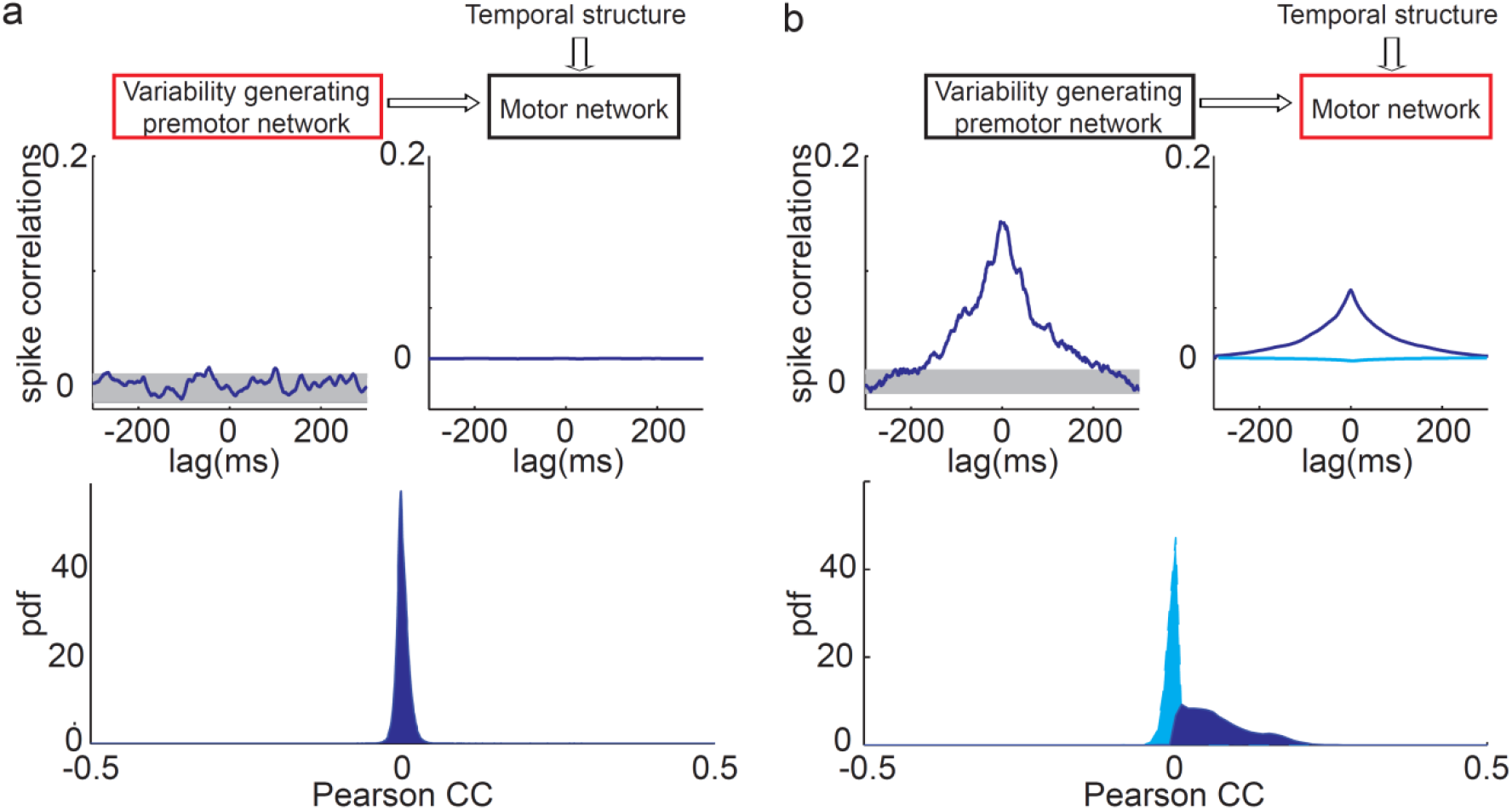
**Correlations in the trial-to-trial neuronal variability increase along the circuit generating motor variability in the model. a-b.** Noise correlations in the model. In the premotor network, Noise crosscorrelations (CCs) are weak. In the motor network, neurons activating the same effector have significant positive correlation coefficients. **a.** Top left: Example of noise CCs across two single-units in the variability-generating premotor network (shaded area: 2.5 SD around the mean). Top right: Population averaged CCs. Noise CCs are almost flat, indicating the absence of significant correlations in the activity of the premotor network. Bottom: Probability density function (pdf) of Pearson correlation coefficients in the premotor network. **b.** Same as in (a), but for neurons in the motor network. Bottom: Conditional probabilities of the Pearson correlations ofneurons in the same functional group(dark-blue; average correlations: ~0.068) and neurons in different groups of (light blue; average correlations: average correlations: ~ -0.0066). Note that the probability of having two neurons in a group and between groups depends on the number of groups and by taking these priors into account the average correlations across all neurons is close to zero (average correlations: ~0.0008; see text and Supplementary Information).

Importantly, the correlations in the motor network are substantial only if the footprint of the recurrent interactions in that network is sufficiently wider than the footprint of the premotor-to-motor projections (Fig.2h; see also Supplementary Information). Indeed, when the recurrent interactions are too local, correlations in the motor network are weak (Fig.2h-i). Thus, temporally irregular *and* spatially correlated patterns of activity naturally emerge from the interplay between topographic feedforward projections from the premotor to the motor network and recurrent interactions within the motor network (Supplementary Fig.10).

The emergence of spatial correlations in the motor network can be intuitively understood as follows. The feedforward (FF) input to a neuron in the motor network consists of one structured component, shared by all neurons belonging to the same functional group, and another one which is unstructured. Since the neurons in the premotor network are firing asynchronously, both components are the sum of a large number of uncorrelated contributions (on average *fK* and (1 − *f*)*K*, respectively; *K* being the average number of synapses per neuron; see Supplementary Fig.3i) and thus their temporal fluctuations are smaller than their temporal average by a factor on the order of 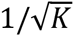 (Supplementary Fig.3f: blue curve). Neural activity in the motor network will be spatially correlated if the amplitude of the fluctuations in the structured component to the network is on the order of the neuronal threshold. However, this implies that the temporally averaged FF input is large. As a consequence, neurons in the motor network will fire regularly at a very high rate, unless the inhibitory recurrent inputs in the motor network compensate for most of its averaged FF input. This compensation will occur naturally for a strongly connected motor network operating in a regime where inhibition balances excitation (see Supplementary Information).

The fluctuations in the component common to all neurons in the same functional group give rise to the correlations in the activity in the motor network on a spatial scale on the order of the size of a group. Moreover, the groups also compete with each other provided that the recurrent interactions extend over a distance larger than the size of a group. As a result, the network dynamics self-organize such that the average instantaneous rates of the excitatory and inhibitory populations are essentially constant in time (Fig.2d). This guarantees that the network operates in the balanced excitation-inhibition regime in a robust manner (see Supplementary Information).

### Emergence of spatiotemporal correlations in the circuit driving vocal variability in songbirds

Songbirds, with their well-identified and segregated circuit devoted to song learning, including a minimal circuit driving song variability (see Introduction), offer an ideal opportunity to test predictions of our theory. In songbirds, LMAN controls the trial-to-trial fluctuations across repetitions of the temporally structured song^8-9^. These fluctuations are important for adapting the song upon perturbations^20-22^. Moreover, anatomical studies in the circuit driving the song indicate that the projections from LMAN to RA are topographically organized, as our model posits for the projections in the premotor-to-motor pathway^23-24^. We therefore hypothesized that song variability stems from essentially uncorrelated fluctuations produced in LMAN, which by virtue of the topographic projections from LMAN to RA induce spatially correlated fluctuations in RA activity. To further test this hypothesis, we extended the two-area circuit considered above to take also into account the temporally structured inputs from nucleus HVC (used as a proper name) into RA neurons^12,25^. To this end, we included an additional feedforward excitation to the motor network in our model, representing the latter input (Fig.3b, see Materials and Methods). The responses of the neurons in the motor network are then locked to this input in a way which is reminiscent to the locking of RA neurons to the song^26-27^ (compare Fig.3a and Fig.3c). However, these responses still exhibit trial-to-trial variability. By analyzing the spatiotemporal patterns of these trial-to-trial fluctuations, we found substantial noise correlations (see Materials and Methods) for neurons in the motor network belonging to the *same group*, but almost none in the upstream premotor network (Fig.4). In the motor network, noise correlations were positive for pairs in the same group. They were typically weaker and negative for pairs in different groups. The averaged correlation over all pairs of excitatory neurons was very small due to the compensation between positive and negative correlations. (Fig.4b and also Supplementary Figure 4). Our model thus predicts a build-up of noise correlations along the circuit generating behavioral variability in singing birds.

To test this prediction, we recorded pairs of LMAN or RA neurons during singing in zebra finches. In LMAN, we found that spike-triggered-average (STA) of the local field potential (LFP), as well as STA of the multi-unit activity are weak (Fig.5b, Supplementary Fig.6, see Materials and Methods). We also found that noise crosscorrelograms are flat (Fig.5c-d) and that correlation coefficients are tightly distributed around zero (Fig.5e) in LMAN. By contrast, in RA neurons display substantial noise correlations during singing, as revealed by the shape of their crosscorrelograms (Fig.5h-i; one-tailed two-sample t-test; p<0.01 for single-units pairs, n=4 pairs in LMAN and n=5 pairs in RA; p<0.001 for single- Vs. multi-units pairs, n=6 pairs in LMAN and n=25 pairs in RA; and p<0.001 for multi-units pairs, n=21 pairs in LMAN and n=21 pairs in RA; see Materials and Methods) and large values of noise correlation coefficients (compare Fig.5j with Fig.5e. The fact that correlations in RA were stronger than in LMAN is consistent with the model prediction, since the recorded units were likely to be located in the same functional group given the small distance between electrodes compared to RA diameter (see also Supplementary Fig.6b). Multi-unit-STA and LFP-STA also consistently display high noise-related activity around the recorded spikes in RA, in contrast to LMAN (compare Fig.5g and 5b and Supplementary Fig.6a). Finally, we found that noise cross-correlations between LFPs recorded from evenly spaced electrodes decreased with the distance between the electrodes and became negative when they were far apart (Supplementary Fig.6b). Therefore, as predicted by our model, noise correlations during singing are strong in RA, while they are extremely weak in LMAN.

**Figure 5.**
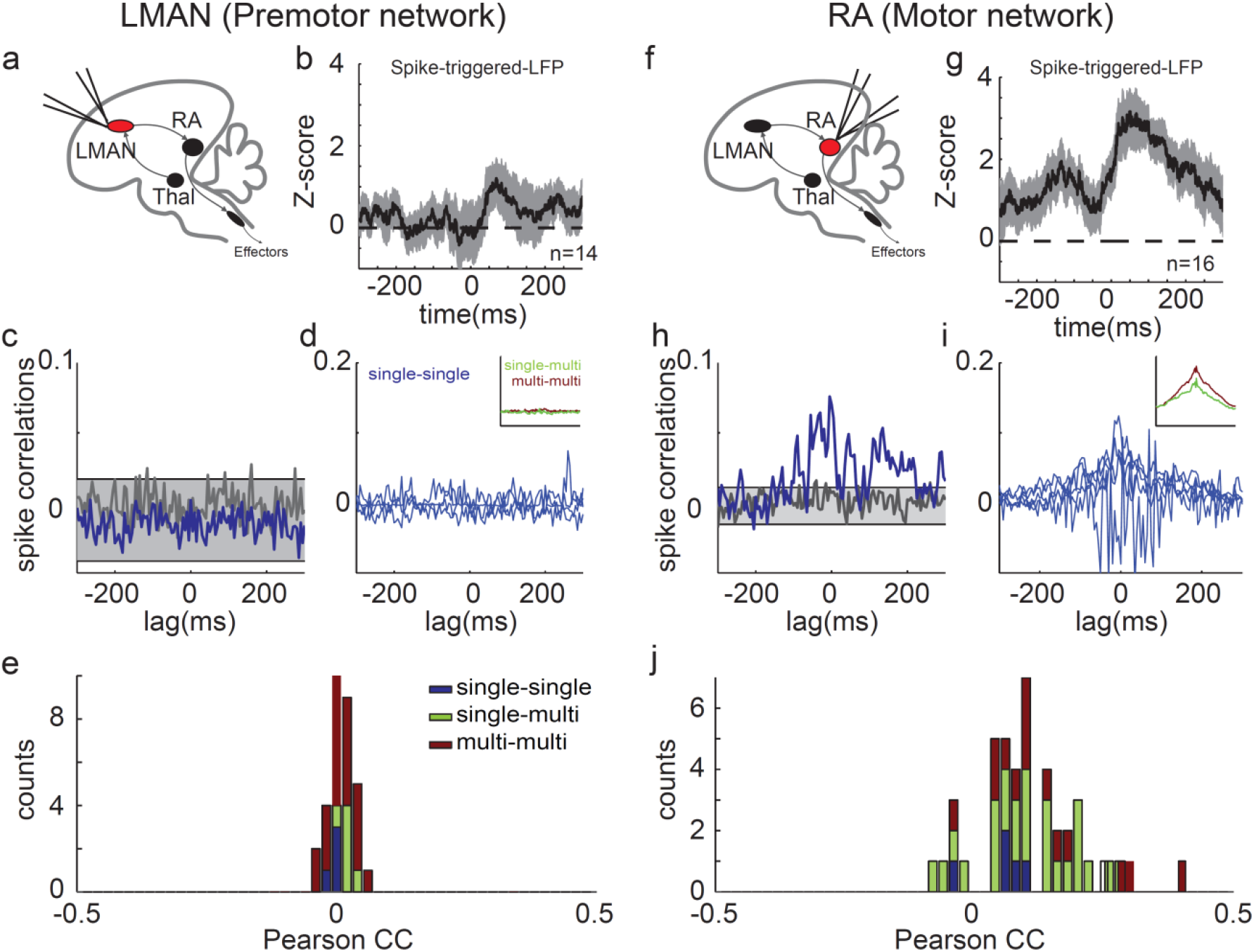
**Correlations in the trial-to-trial neuronal variability increase along the circuit generating motor variability in singing birds.** Experimental recordings in zebra finches during singing. Noise correlations are weak in LMAN but substantial in RA. **a**.Area of recordings. **b.**Spike-triggered average (STA) of the noise-LFP during singing in LMAN (mean±s.e.m.). The motif-average LFP was subtracted from the LFP signal and the STA of this residual LFP was then computed separately for each single unit recording (see Materials and Methods). **c.** Noise CCs of two single-units recorded simultaneously in LMAN during singing. The mean motif-related activity was subtracted from the instantaneous firing rate during singing and correlation analysis was performed on the residual trial-to-trial fluctuating signal (noise correlations, see Materials and Methods). Noise CCs are flat after a random permutation of the spikes (gray trace, shaded area: 2.5 SD around the mean). **d.** Single-unit pairs crosscorrelograms (blue) and average crosscorrelograms (inset) of single- vs. multi-units pairs (green) and pairs of multi-units (red) recorded from different electrodes in LMAN. **e.** Distribution of Pearson correlation coefficients in LMAN.**f-j.**Same as (a-e), but for neurons recorded in RA. In contrast to LMAN, RA neurons exhibit significant pairwise correlations. Crosscorrelograms are broad and their integrals are significantly larger in RA than in LMAN (see Materials and Methods and Results for statistical tests), reflecting the slow co-fluctuations in the activity of the simultaneously recorded units.

Our electrophysiological recordings also reveal that the decays of the autocorrelations and of the crosscorrelations of the spiking activity last for hundred of milliseconds in RA neurons (Fig.6b-c;Fig.5h-i), and that these decays are substantially faster in LMAN (Fig.6a,c; two-sample t-test, p<0.01 for n=10 single-units in LMAN and n=14 single units in RA). What is the source of the relatively slow decorrelations in the activity of RA neurons? In our theory, *synchronous* temporal fluctuations in RA activity will be slow if the shared feedforward drive of the neurons in the motor network slowly fluctuates (see also Supplementary Information and Supplementary Fig.7). If the synaptic dynamics in the premotor-to-motor pathway are slow, they give rise to auto and cross-correlograms in the motor network which can be as broad as in the data (compare Fig.6a-c with Fig.6d-f and Fig.5h-i with Fig.4b). This result suggests that the observed slowness of the fluctuations in RA activity stems from a low pass filtering of the fast fluctuations of LMAN outputs due to the large proportion of NMDA receptors in the LMAN-to-RA projections^28-30^.

**Figure 6.**
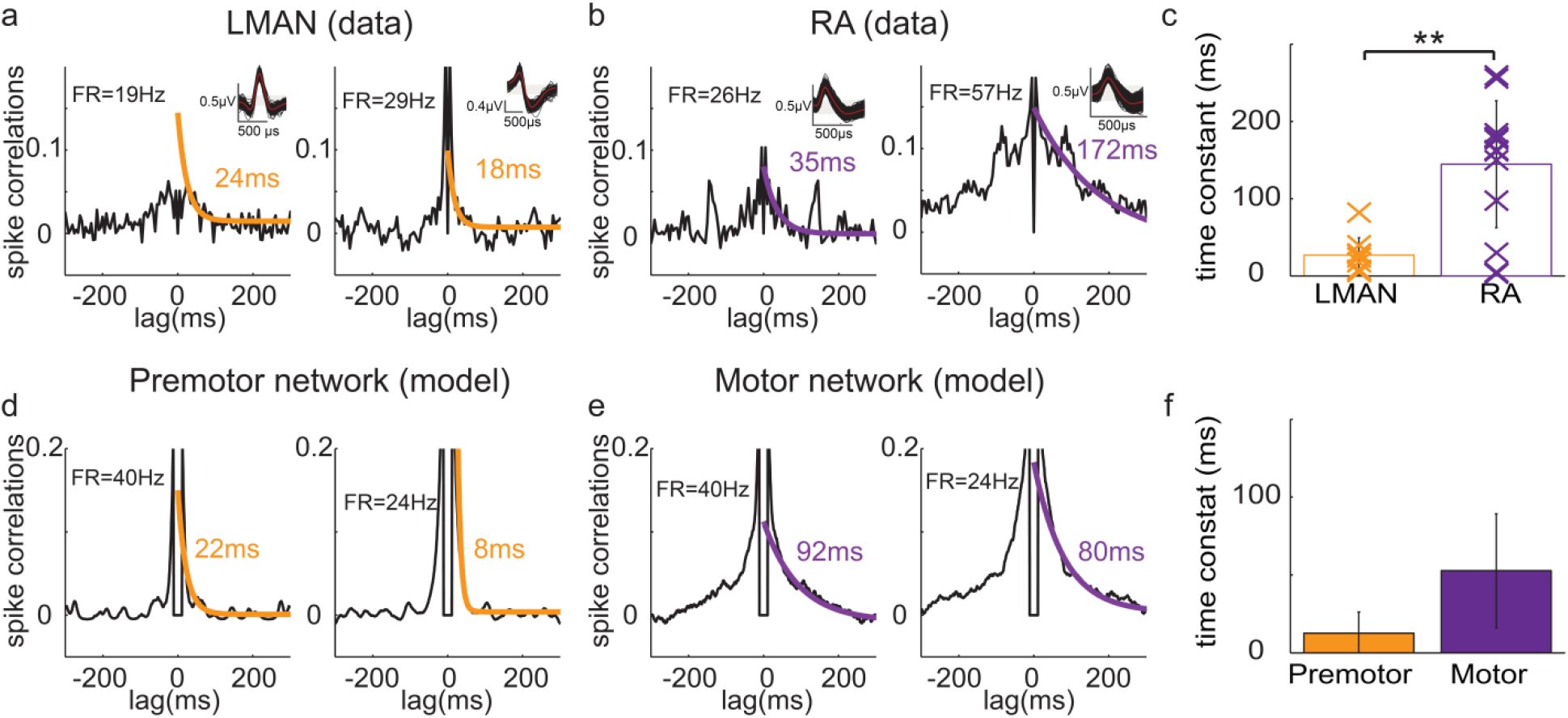
**Temporal fluctuations slow down along the circuits generating motor variability (data+model). a.-b.** Decorrelation time in the activity of neurons in LMAN or RA during singing in zebra finches. **a.** Two examples of noise autocorrelations (ACs, see Materials and Methods) for neurons recorded in LMAN (simultaneous recordings, CCs plotted in Fig.5c). Inset: superimposed spike shapes (red: average trace). FR: average singing-related firing rate during song. **b.** Same as (a) but for two RA neurons (simultaneous recordings, CCs plotted in Fig.5h). The ACs are fitted to a decaying exponential (Orange in a and purple in b; Time constant is indicated in the panels). **c.** ACs are much broader in RA than in LMAN (single units and mean+s.d.).**d-f.** The same as in (a-c) but in the model (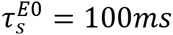, see Materials and Methods). **d.** ACs for neurons in the premotor network. **e.** ACs for neurons in the motor network.**f**.ACs in the premotor network decay faster than in the motor network (mean+s.d.).

### The statistics of vocal variability in juvenile learners are similar across species as predicted by the model

Juvenile songbirds produce babbling-like vocalizations which are not stereotyped and highly variable^31,10^. At this early developmental stage, the inputs from HVC to RA are not yet functional^32^ and the song is mostly driven by LMAN-RA circuit^10,12^. We therefore asked whether the neuronal circuit depicted in Fig.2a can drive behavioral variability with statistics similar to those observed during the babbling stage of juvenile birds. To this end, we combined the circuit with a mechanical model of the vocal production organ^33^. To characterize the statistics of the output signal of the model we computed the distribution of gesture durations (vocal elements) and the autocovariance of the envelope signal (ACE; see Materials and Methods), which quantifies high-order correlations between consecutive gestures and inter-gesture-intervals. The gestures durations had an exponential distribution in the model (Fig.7a, bottom left; see Materials and Methods). As for the ACE (Fig.7a, bottom right), it monotonically decays over a duration of several tens to hundreds of milliseconds. Quantitatively, the decay time constant of the ACE and the scale parameter of gesture duration distributions depend on the synaptic time constant of the premotor-to-motor projections (compare the blue and red line in Fig.7a).

**Figure 7.**
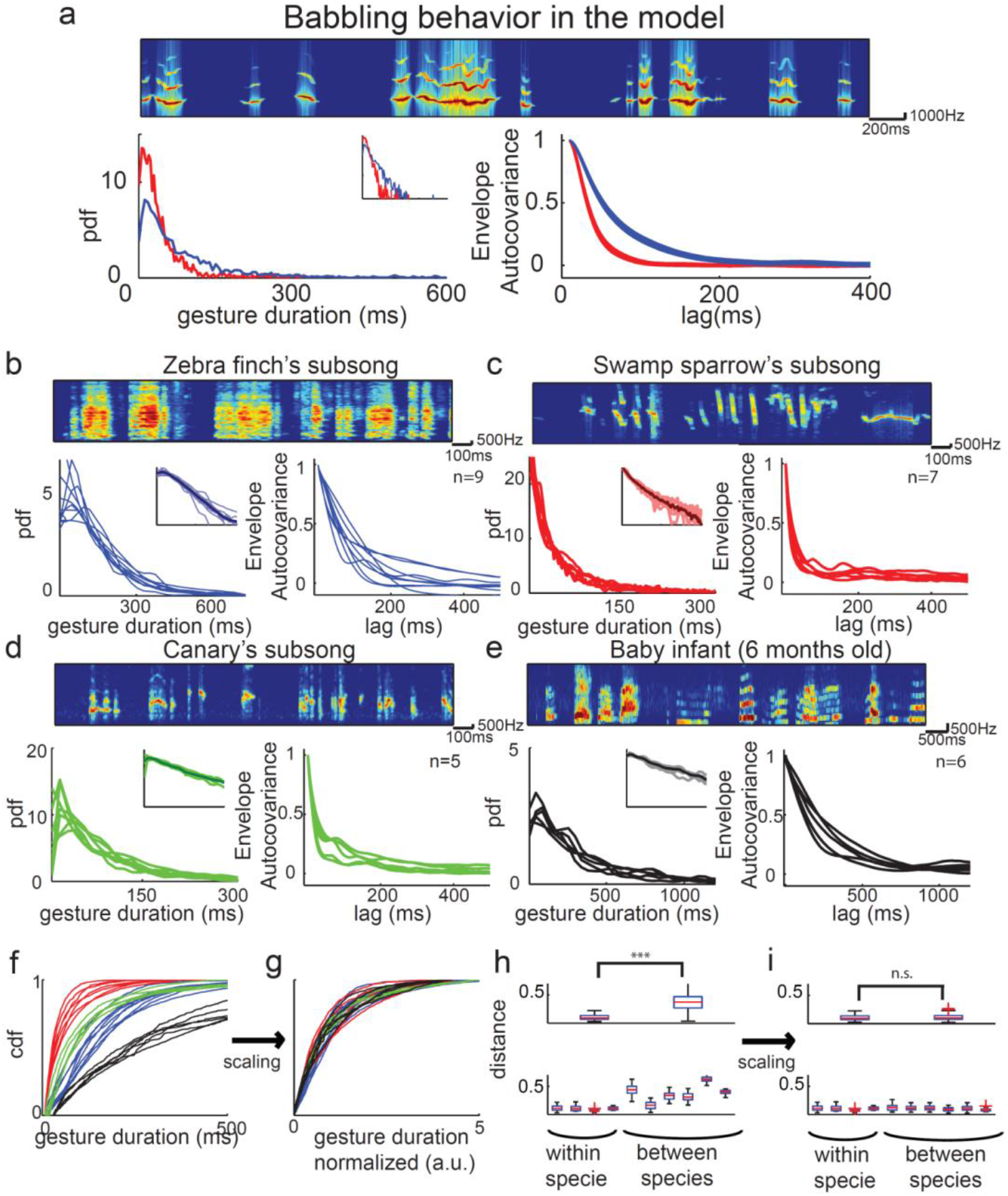
**Babbling statistics are similar across different species of songbirds, human infants and in the model.a.** Statistics of the babbling behavior generated by the model circuit depicted in Fig2. a-e when coupled to a minimal model of the vocal organ (see Materials and Methods). Top: spectrogram of the vocal output signal 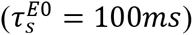. Bottom: probability density function (pdf) of vocal gesture durations (left) and averaged autocovariance of the envelope (ACE; right). Inset: distribution of gesture durations when the Y-axis is in log-scale. The distribution of gesture durations is well approximated by an exponential with a ‘scale parameter’, 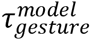 (see Material and Methods). ACE decorrelates over a time duration of 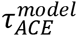. Slow synaptic dynamics in the premotor-to-motor projections (red: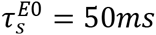; blue: 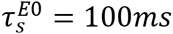) results in slowly fluctuating vocal output (red: 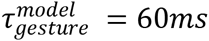 and 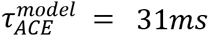; blue: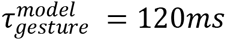 and 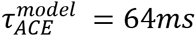). **b-i.** Statistics of the babbling behavior in four species of vocal learners(ages of the subjects (‘babbling period’)are given in Material and Methods). Blue: Zebra finches (Zf); Red: Swamp sparrows (Sw); Green: Canaries (Ca); Black: Human infants (Bab).Different lines of the same color correspond to different subjects from the same species. **b-e.** Same as in a, but for the Zf (b: compare to the blue line in (a)), Sw (c: compare to the red line in (a)), Ca (d) and Bab (e). Gesture duration distributions lack any clear peak and are well fit with exponential decaying function with scale parameters (mean±s.e.m): 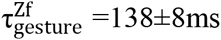; 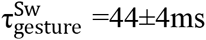; 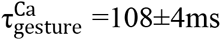; 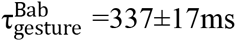. The ACE decay time is specie-dependent: 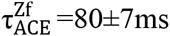; 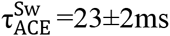; 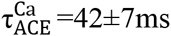; 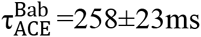**.f-g.** Cumulative distribution functions (cdf) of gesture duration for the four species before (f) and after (g) normalizing the gesture durations by 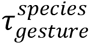. **H.** Top: Interspecies differences in cdfs are much smaller than intraspecies differences (Kolmogorov-Smirnov statistic as a distance measure between cdfs). Bottom: Differences of cdfs in pairs of learners within (left to right: Zf-Zf, Sw-Sw,Ca-Ca, Bab-Bab) and between species (left to right: Zf-Sw, Zf-Ca, Zf-Bab, Sw-Ca, Sw-Bab, Ca-Bab). **i.** Most of the *interspecies* differences in (h) are accounted for by normalizing the gesture durations to the scale parameter of the exponential fit of their distributions (see Results and Material and Methods for statistical comparisons).

To what extent do these statistics depend on the details of the model architecture and connectivity, the neuronal dynamics, and the nonlinearities in the input-output transduction by the effectors? Fluctuations in the feedforward input to the neurons in the motor network in our circuit consist of many uncorrelated fluctuating contributions and their statistics are thus close to Gaussian (Supplementary Fig.8a). This is the case also for the net (feedforward+recurrent) input to these neurons (Supplementary Fig.8b). Hence, the fluctuating activity of the neurons in the motor network can be approximately described as a wideband Gaussian process that is rectified (Supplementary Fig8.b-d), resulting in the tendency of the neurons to fire bursts of spikes with an approximately exponential distribution of durations (Supplementary Fig.8f). Moreover, because neurons in the motor network are correlated, the temporal statistics of the input to an effector and of the neurons activity are similar. While the babbling behavior generated by the circuit is a complex transformation^34^ of the inputs to the effectors, the ACE or the distribution of gesture durations (but not finer structures of the gestures) are expected to be qualitatively independent on the details of this transformation. For example, for rectified power-law transformations the distribution of gesture durations is close to exponential (Supplementary Fig.8g) and the ACE barely depends on the non-linearity (Supplementary Fig.8e). This is also true when combining the circuit with a mechanical model of the vocal production organ^33^. Therefore, these features reflect universal statistical properties of the circuit dynamics and are, to a large extent, insensitive to the circuit parameters and to the details of the transformation from the input to the effectors to the vocal behavior.

Given the universality of the statistics of the features analyzed above in our model, we then compared the “babbling behavior” generated by the model with babbling behavior in different vocal learners. To this end, we analyzed babbling vocalizations of juveniles from three different songbird species with completely different adult repertoires (zebra finches: single song of 3-8 syllables per individual; swamp sparrow: 2-5 stereotyped songs per individual gathering 5-10 syllable types; canaries: complex song sequences based on a repertoire of 20-40 syllables per individual), as well as vocalizations of 5-6 months old human infants (adult repertoire: complex sentences based on 10-100 phonemes grouped in >10 000 words).

Remarkably, we found that the statistics of the vocalizations produced during the early period of babbling (but not later in development, Supplementary Fig.9a-c,f) had a large degree of similarity in the four species we analyzed. In all four species, as in the model, the distribution of vocal gesture durations could be well fitted with a single exponential (Fig7b-e, left and insets; see Materials and Methods, see also (10,35)). In addition, the ACE lacks a clear temporal structure in all babbling vocalizations. The scale parameter of the gesture duration distributions, as well as the decorrelation times of ACE (i.e., the typical time constant of the ACE) varied across individuals and species from several tens to a few hundreds of milliseconds (Fig.7b-e). However, as the distributions were close to exponential and variability within species was small (Fig.7f,h), interspecies and intraspecies differences in gesture duration distributions became comparable after normalizing each individual distribution by its *species-averaged* scale parameter (Fig.7g-i; two-sample t-test, P=0.92: only 9% of the total variance among distributions was attributed to species differences compared to 87% before rescaling; 1-way ANOVA). A similar uniformity was observed in the inter-gesture-interval distributions after normalization by the species average (Supplementary Fig.9d.; P=0.23, only 10% of total variance among distributions was attributed to species differences compared to 66% before normalization; 1-way ANOVA). Interspecies variations in ACE were mostly due to differences in the species-averaged decorrelation time (P=0.16; only 17% of the total variance among ACEs was attributed to species differences, compared to 81% before normalization; 1-way ANOVA). Finally, correlations between consecutive gestures and inter-gestures were small and comparable among species (Supplementary Fig.9e). Together, these results show that in the four species we studied the statistics of the babbling-like vocalizations are very similar and can be naturally accounted by our minimal circuit.

## DISCUSSION

Our paper addresses the extent to which the intrinsic temporal irregularity of neuronal activity in the central nervous system (CNS) can drive motor variability. This is a fundamental non-trivial question since, as to impact the behavior, patterns of activity generated in the CNS must also be *spatially* correlated (i.e., correlated across neurons). Although the emergence of asynchronous irregular activity in recurrent networks is well understood^13-16^, much less is known regarding the possible mechanisms giving rise to irregular spiking in which fluctuations of the activity are both temporally irregular *and* correlated across neurons. As a matter of fact, in virtually all network models of irregular spiking previously investigated, the activity is either asynchronous^16^ or the synchronous component of the temporal fluctuations in neuronal spiking is strongly rhythmic^36^.

In particular, previous theoretical studies^16,37^, concluded that correlations should be very weak in strongly recurrent cortical circuits (on the order of 1/N, where N is the network size). However, these studies assumed a completely random connectivity, without structure (with an Erdös-Renyi graph). Here we showed that substantial correlations emerge naturally in a circuit with topography. With such an architecture, the dynamics self-organize in groups of neurons that are positively correlated within a group but negatively correlated between groups. In this spatial pattern of correlations, the balance between excitation and inhibition is maintained over the whole network. As a result, the circuit can eventually produce robust variable behavior with ‘universal’ statistics. In fact, we showed that this mechanism does not require *any fine-tuning of parameters.* In particular, it is robust to the number of neurons, the average number of connections, as well as to the connectivity in the topographic pathway to the effectors (and the number of neurons projecting to an effector; see also Supplementary Information).

In songbirds, the organization of the LMAN-to-RA pathway becomes clearly topographic during the early sensory period of song learning^31^. Thus, it is already present when juveniles start to babble (35-40 days post hatch, DPH). Neurons in RA also send topographic projections to the hypoglossal nucleus (nXII) as well as to the respiratory motor nuclei^24^. The projections of the hypoglossal nucleus to syringeal muscles are also topographic^38^. Thus the pathway from RA to syringeal muscles (and likely similarly to respiratory muscles) is topographic, as required by our mechanism. Applied to the LMAN-RA circuit, this mechanism predicts that noise correlations are weak in LMAN but substantial in RA. We reported experimental evidence in line with this prediction in the adult zebra finch.

In juvenile and adult zebra finches, the inputs from LMAN to RA are dominated by NMDA receptors with slow kinetics of time constant on the order of ~100ms^28-30^. Moreover, recurrent excitation in LMAN is largely dominated by NMDA receptors and the kinetics of these receptors is faster in adults than in young juveniles^39^, with typical time constants of ~30ms in adults and ~120ms in juveniles^40^. Therefore, slow synapses in LMAN as well in LMAN to RA projections can underlie the relative slowness of the dynamics of the babbling behavior we reported in juvenile finches (se also Supplementary Fig.7 and Supplementary Information). In agreement with this view, localized mild cooling of LMAN in zebra finches results in an increase in the time constant of the exponential gesture distribution during babbling-like behavior and in a longer tail in the distribution in older juveniles^35^.

Our behavioral data show substantial differences in the time scale of the babbling behavior between zebra finches, canaries, swamp sparrows and humans. Our model suggests that this may be due to differences in the kinetics of NMDA receptors in thesespecies.Revealing a direct correlation between these differences and NMDA receptors kinetics requires data on the latter. To the best of our knowledge, there is no such data available for canaries, swamp sparrows or human infants. However, the range spanned by the babbling time scales in our behavioral data is compatible with the diversity of kinetics reported in NMDA receptors of different sub-unit composition^41,42^.

In adult subjects, motor variability is expressed as fluctuations around a stereotyped motor pattern, which despite their relatively small amplitude, can contribute significantly to motor learning^2^. At early stage of development young animals, as well as human infants, produce spontaneous exploratory gestures referred to as “motor babbling” that do not rely on any stereotyped or goal-oriented movement, and rather appear to express pure motor variability^10,43-44^. Such exploratory movements may allow the self-organization^45^and the adaptation of sensory-motor networks through correlation-based (Hebbian learning) and reinforcement learning mechanisms^1,46-48^. These mechanisms posit that synaptic neural correlates of exploratory behavior must persist for tens of milliseconds in the learning circuit. Our work suggests that the wide presence of NMDA receptors in the LMAN-to-RA projections^29-30^ is a key component in the emergence of such *eligibility trace* in the overall dynamics of the circuit which generates behavioral variability in birds.

To conclude, we showed that a circuit comprising strongly recurrent neural networks, which is organized in a topographic manner, is capable of driving variable motor behaviors. This mechanism relies on only a few architectural constraints and is thus likely to be a general operating principle by which the brain acquires motor skills and adapt behavior in a changing environment.

## Acknowledgments

We thank Shaul Druckmann, Robert Gütig, Carole Levenes, Gianluigi Mongillo, Richard Mooney, Israel Nelken, David Perkel, Nicholas Priebe, Frederic Theunissen and Tatyana Sharpee for comments on the manuscript. We thank Catherine Del-Negro and Karine Martel for sharing with us some of their behavioral data and the parents of the human infants for recordings their infants. Work conducted in the framework of the France Israel Laboratory of Neuroscience. Grants: ANR SONGLEARN, ANR. BALWM, Marie Curie IRG-NMVLRBG, France-Israel High Council for Science and technology, LIA FILN.

R.D. and A.L. performed the behavioral data analysis. S.P. provided the sparrows data. W.E.W. and A.L. performed the electrophysiological experiments. R.D., W.E.W. and A.L. performed the electrophysiological data analysis. R.D. and D.H. designed the theory and performed the simulations. R.D., A.L. and D.H. designed the study and wrote the manuscript.

## ONLINE METHODS

### Sound Recordings

#### Subjects

Seven human infants (3 males and 4 females) were recorded in their natural environment. Their parents gave written informed consent for participation in this study. Nine zebra finches and five canaries were obtained from our breeding facilities (Paris Descartes and Paris Sud Universities). Seven swamp sparrows were collected as nestlings and hand-reared in the laboratory (see ref. (50) for details). Birds were housed under natural light/dark conditions and provided with food and water ad libitum. Animal care and experiments were performed in accordance with European directives (86/609/CEE and 2010-63-UE) and the French legislation. Experiments were approved by Paris Descartes University ethics committee.

#### Human infants

We recorded spontaneous vocalizations in six infants in their natural environment starting from 5 to 7 months after birth(denoted as the: ‘babbling period’). The parents were instructed to place a recorder (digital dictation machine with stereo microphone, ICD-PX333M SONY) near the baby’s head for ~30 min at least 5 days a week for several weeks (4 to 20 weeks). The data presented include babies for which vocalizations were collected from at least 20 days during this recording period. Additionally, one 10 month old infant was recorded for repetitive babbling.

#### Zebra finches

Juvenile zebra finches were raised in single cages with their parents and siblings. At age 26–41 DPH(day-post-hatch, ‘babbling period’), 9 male zebra finches were removed and placed in custom-made sound isolation chambers. Vocalizations were recorded for 10-30 days continuously with Sound Analysis^51^, which was configured to ensure that recordings were triggered on all quiet vocalizations of young birds. Five of the nine birds were continuously recorded until song crystallization (~3 months), with episodic access to their father.

#### Swamp sparrows

Seven swamp sparrow males were recorded in individual sound isolation chambers (Industrial Acoustics AC-1) once per week, starting in February of their first year when they were about 250 DPH. The onset of song development was first detected at 262-296DPH(‘babbling period’), and recording continued up to 366-386 DPH, when the males were singing crystallized adult song. Subsong was sampled for 30 minutes (Marantz PMD221 cassette tape recorder, Realistic Omni-directional microphone, Yamaha Mike to Line Amplifier). An automated system was introduced to detect and record song during late-plastic and crystallized song using a voice activated switch (modified UherAkusomat) and a Digital Delay System (Digitech).

#### Canaries

Juvenile canaries were raised in our breeding facility at Paris Sud University, in single cages with their parents and siblings. At age 75–150DPH(‘babbling period’), as they started to produce their first vocalizations, five male canaries were removed and placed in custom-made sound isolation chambers. Vocalizations were recorded continuously for 3 months (September to December) during the fall following their birth with Sound Analysis Pro, which was configured to ensure that recordings were triggered on all quiet vocalizations of young birds. Four of the five birds were also recorded 3 months later (early spring)for 5-10 more days.

### Sound analysis

#### Vocalizations

Songs and infant vocalizations were manually sorted. For subsongs we took the first recorded song vocalizations of the bird. Recordings were from one day of vocalizations, except for zebra finches, where in some individuals subsongs from 1-3 recording days were combined to get enough gestures.

#### Spectrograms

Spectrograms were estimated using the multitaper method with 2 slepian tapers.

#### Envelope signal

We extracted the envelope of the signal (termed also “amplitude” in the literature) by band passing the sound signal in the frequency ranges of the vocalizations (from 800Hz and up to 4000-10000Hz, depending on the species, with order-80 linear-phase finite-duration impulse response, FIR, filter), taking the absolute value of the signal and low passing it at 1-200Hz with a linear filter of order-200 linear-phase filter FIR.

#### Averaged Auto-Correlations of Envelope (ACE)

The autocovariance of the envelope signal was estimated for each recording and then normalized to the zero lag. The ACE signal was then estimated by averaging this signal over one day of recording sessions.

#### Gesture and Inter-Gesture segmentation

We used a local method for gesture and inter-gesture detection. We calculated the peaks of the derivative of the log-envelope signal (after band passing the signal; see above) which was smoothed by a 5-30 ms sliding window, depending on the noise level of the signal and the species (using fpeaks in Matlab and a filter of (-1 0 1)) and defined sound onsets and offsets as the closest points to these crossings. We defined the threshold for each file by the x percentile of the peaks, where x was in the range of 85-98. The percentile threshold, x, as well as other relevant parameters for segmentation were fixed after manually examining a subset of the data for each recording day. When the signal was too noisy to use the local method (mainly for infant babbling and several swamp sparrows) we used a global threshold: for each recording, we calculated a sound threshold by fitting a two-Gaussian mixture model (corresponding to noise and sound) using an expectation-maximization algorithm for the log-envelope signal. We then detected crossings of this threshold and defined sound onsets and offsets as the closest points to these crossings where the envelope deviated from the noise by 4 standard deviations. Using a global method instead of a local one on all the juvenile recordings yielded similar results, however we preferred to use the local method whenever possible. For both methods, sounds separated by a duration of <7ms of silence were merged into a single gesture, and segments of overly long (Zf: >800ms; Sw:>400ms; Ca: >900ms; Bab: >3500ms) or short durations (Zf, Sw,Ca:7ms; Bab:<30ms) were eliminated.

#### Fitting exponential decay

We fit an exponential function to the gesture duration distribution using maximum-likelihood estimation on a finite interval^35^ (based on a median of 1870 gestures per day for a songbird and 700 for about one month of human infant recordings; interval duration: Zf: 50-800ms; Sw:10-200ms; Ca: 50-600ms; Bab: 50-1500ms). Distributions that were well fit by the exponential function usually had a high goodness-of-fit metric^35^ (adjusted-R^2^>0.7 and Lilliefores statistic<3.5). To extract decorrelation timescales from the ACE we fit an exponential decay to the ACE.

### Electrophysiology

#### Microdrive implantation

For single and multi-unit recordings, a custom-built motorized microdrive (RP Metrix) was modified to accept 2-4 tungsten microelectrodes (8-20 MΩ, FHC), as well as lateral positioner. It was implanted in LMAN or RA as follows: Young adult male zebra finches (<180 DPH, 3 for nucleus LMAN and 2 for nucleus RA) were anesthetized with 5% isoflurane (induction) and placed in a stereotaxic apparatus with a head angle of 30-50°(for LMAN implantation) or -5° (for RA). Anesthesia was maintained with 0.5-1% isoflurane for the duration of the surgery. LMAN/RA was located using previously established stereotaxic coordinates and identified based on its characteristic neural activity patterns. The electrodes and exposed brain were surrounded with Kwik-Cast (WPI), and the microdrive was secured to the skull using dental cement (Superbond, Phymep). A silver wire implanted under the skull acted as a ground, and a low impedance fixed tungsten electrode served as the reference. We also conducted LFP recordings in nucleus RA, using 3x3 micro-electrode arrays (AlphaOmega) with an impedance of 1 to 2 MOhm. Electrode arrays were implanted in young male zebra finches under isoflurane anesthesia (as specified above), relying on the recorded signals to locate nucleus RA, and then fixed onto the skull using dental cement (Superbond, Phymep). A silver wire implanted under the skull was used as a ground, and one of the contact points of the array served as the reference. In both types of experiments, subjects were allowed to recover and habituate to the weight of the recording apparatus for a few days. They were then transferred to the recording cage and connected through a commercial tether and head stage (Neuralynx or AlphaOmega) and the implanted microdrive to a mercury commutator on the roof of the cage (Dragonfly systems). An elastic thread built into the tether helped to support the weight of the implant. Subjects remained tethered both during and between experiments.

#### Chronic recordings

Neural signals and vocalizations were collected using a commercial head stage and acquisition system (Neuralynx or AlphaOmega). Signals were amplified, digitized and filtered either below 300 Hz (LFP signal) or between 300Hz and 30kHz (spike signal).

#### Data analysis

Spike signals were analyzed using Spike2 software (Cambridge Electronic Design) and custom-written software in Matlab (MathWorks). Single and multi-unit signals were isolated using Spike2, and spike times were then exported to Matlab. Motif onset times were extracted from sound recordings using custom programs. We calculated the auto-correlograms (AC; 5s window, 5ms bin) of single spike trains and cross-correlograms (CC; 5s window, 5ms bin) of all pairs of spike trains recorded simultaneously. Activity was first aligned to motif onsets and then average over all motifs produced during each recording session (PSTH analysis), using a time window limited to the duration of a single song motif, and 5 ms bins. To eliminate temporal variability due to fluctuations in the duration of single syllables, spike trains were aligned and stretched using piecewise linear time warping with each syllable onset as a time reference^27^. Signal correlations measure the similarity of the activity of two neurons during singing. Noise correlations, on the other hand, is a measure of the similarity of the trial-to-trial variability (around the motif-related PSTH) of two neurons. We computed the spike counts in 5 ms bins during song production and subtracted the mean motif-related PSTH of a song motif for each neuron for all motifs produced. For each pair of simultaneously recorded neurons (by definition, the noise correlations can only be calculated for simultaneously recorded neurons), we computed the noise correlations by calculating the correlation coefficient of these two vectors. To compare the shapes of the cross-correlograms from units recorded in two considered brain nuclei (LMAN and RA), we measured the mean deviation from zero in a cross-correlogram according to the following procedure. The absolute value of the cross-correlation function was first averaged over the [-50ms +50ms] window (appropriate for the behavioral output given the integration time-constant). The absolute value of the auto-correlation functions of the two corresponding units was also averaged over the same time window. The average absolute cross-correlation was then normalized to the square root of the product of the average absolute autocorrelation functions to provide the numbers used in the statistical tests in the main text. This equation is given by:

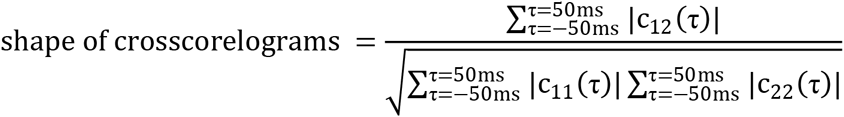

wherec_12_ (τ) is the cross-correlogram across the two neurons at time lag τ, and c_11_ (τ) and c_22_ (τ) are the autocorrelations of the two neurons.

LFP signals were aligned to motif onsets and averaged over all motif renditions produced during a recording session. To calculate the “noise Spike-Triggered Average LFP”, we first subtracted motif-aligned average LFP to LFP signals recorded during each single motif. We then computed the average of the residual-LFP signals cut in 600ms window around each spike over all spikes produced during all motif renditions in the recording session. Additionally, we computed noise Spike-Triggered Average of the envelope of the background multi-unit spiking activity present behind single-unit recordings. To this end, we first removed spike shapes from the sorted single-unit from raw spiking signal, rectified the leftover background signal and convolved it with a 5ms wide Gaussian function. A similar treatment as for the LFP was applied to this background multi-unit envelop to get its noise Spike-Triggered Average.

To quantify the possible effect of bad or partial time-warping on the level of correlation in our data set, we first compared the level correlation in our data set before and after time warping, and then also incorporated artificially wrong syllable timing before the time-warping was applied. To this end, we added variable jitter (from 2 to 500ms) to the syllable onset times, and then time-warped the spike times of the two units according to this jittered syllable timing. This manipulation served to artificially introduce a strong misalignment of the spiking activity with real singing behavior, with a common time jitter for both units, without modifying the average activity of the neurons in a given trial.

### Statistics

Numerical values are given as mean ±SD, unless stated otherwise. Whenever using a statistical test, we report the type of test applied and the associated *p* value (probability of observing the given result, or one more extreme, by chance if the null hypothesis is true).

### Histology

After the last recording session, subjects were killed by intramuscular injection of sodium pentobarbital (Nembutal) and perfused transcardially with 0.9% saline followed by 4% paraformaldehyde as fixative. The brain was then removed, postfixed in 4% paraformaldehyde for 24 h, and cryoprotected in 30% sucrose. Sections (60 *µ*m thick) were then cut in the parasagittal plane on a freezing microtome and processed for histological examination to verify the location of the recording electrodes. Tissue was Nissl stained to visualize the electrode tracks.

### Spiking model

Our model consists of two large recurrent networks, both comprising *N*_*E*_ excitatory (E) and *N*_*I*_ inhibitory (I) neurons. For simplicity we take *N*_*E*_ = *N*_*I*_ = *N*. These two networks represent a premotor and a motor network. The premotor network projects in a feedforward manner to the motor network, and the latter activates a small number of effectors, consistent with the songbird anatomy^10,23-24,38^.

#### Single neuron and synaptic dynamics

All the neurons in the circuit are modeled as leaky-integrate-and-fire (LIF) units. The sub-threshold dynamics of the membrane potential, V_i,α_(t), of neuron i in population α (i = 1, …, N_α_; α = E, I) obey:

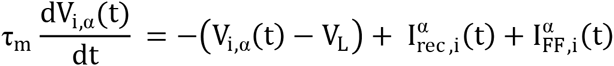

where τ_m_ is the neuron membrane time constant, 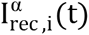 is the recurrent input into neuron (i, α), due to its interactions with other neurons in the same network (premotor or motor), 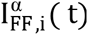 is the total feedforward input into that neuron. V_L_ is the reversal potential of the leak current (taken to be *V*_*L*_ = −60mV).

These subthreshold dynamics are supplemented by a reset condition: if at t = t_iα_ the membrane potential of neuron (i, α) crosses the threshold, 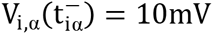, the neuron fires an action potential and the voltage reset to 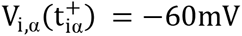.

We model all synaptic inputs as pure currents. The total current into neuron (i,α) due to its recurrent interactions yields:

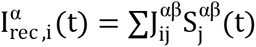

where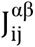 is the strength of the connection from presynaptic neuron (j, β) with neuron (i, α), and 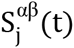 are the synaptic variables, which follow the dynamics:

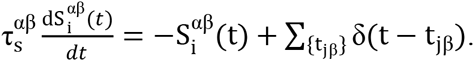

Here 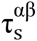 is the synaptic time constant (assumed to depend solely on the nature -excitatory or inhibitory– of the pre and postsynaptic neuron) and the sum is over all spikes emitted at times t_jβ_ < *t*.

#### Recurrent architecture

The recurrent connectivity of the E and I populations in the premotor network is random (Erdös-Renyi graph). In each network, the connectivity matrix, C^αβ^, between presynaptic population β and postsynaptic population α is therefore a random N xN matrix such that 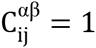 with probability K /N and zero otherwise, where K is the average number of inputs a neuron receives from population β. We assume that the strength of the synapses depends solely on these populations yielding: 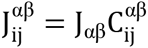 where J_α*E*_> 0 (excitation) and J_α*I*_ < 0 (inhibition). When comparing the dynamics of networks with different connectivity we follow the prescription^13,48^:

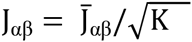

where the parameters, 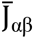, are of order unity and can be different for the premotor and motor networks.

#### Distance-dependent recurrent architecture in the motor network

In the motor network the connectivity is random with probability which depends on the distance between the neurons. The probability of connections between two neurons is 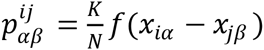, where 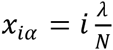 is the location of neuron i=1, …, N in population *α*, and

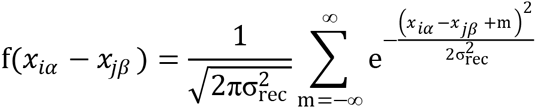

where*σ*_*rec*_ is the footprint of the recurrent interactions. Here we have assumed for simplicity that the motor network is one dimensional, of size *λ* and periodic boundary conditions. For large values of *σ*_*rec*_ distant neurons are as likely to be connected as close ones, while for small values of *σ*_*rec*_ only neurons which are close have a significant probability to be connected (Fig.2h). In most of the results depicted in the paper we assume that the recurrent interactions in the motor network have a wide footprint (*σ*_*rec*_ → ∞), except for Fig.2h-i, where we investigate how the results depend on the value of *σ*_*rec*_.

#### Feed-Forward architecture

The premotor network receives external feedforward (FF) inputs, which in the context of the songbird system represent the thalamic (the medial part of the dorsolateral nucleus of the anterior thalamus, DLM) inputs that may tonically activate LMAN during song. The total number of FF inputs to a premotor neuron is modeled as a constant drive^13,52^, 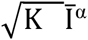. Similarly, the motor network receives a FF input from outside the circuit that we model as a constant drive. Importantly, the motor network also receives FF projections from the premotor area which exhibit a topographic organization. To implement this key feature of the architecture of our model we divide the excitatory population in the motor network into Dstatistically equivalent functional groups (N /D neurons in each group). For each group, we choose a set of *f K* neurons (set *P*_*l*_) in the premotor network projecting to in neurons in the group. For each neuron in the group, additional inputs are chosen by drawing randomly from the premotor network with probability (1 − *f*)*K* /*N*. Each neuron in a group therefore receives on average *K* projections from the premotor network. Changing f allowed us to easily manipulate the total amount of correlations in the FF inputs by changing the parameter, *f*, keeping the total average number, *K*, of premotor inputs per neurons fixed. For *f* = 1 all the projections are topographic whereas if *f* = 0 they are completely random. The total FF premotor input into an excitatory neuron i (*i* = 1, …, N) in group *l*(= 1, …, D) in the motor network is therefore modeled as:

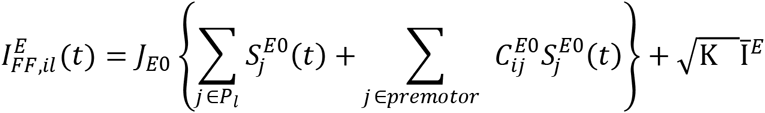

In this architecture, the probability that two neurons in group 1 share premotor inputs is *f* + *O*(*K*^2^/*N*^2^). Note that if *K* is too large it will be impossible to have different shared inputs for each group in the motor network. However, this does not happen with the model parameters in the simulations described in the paper since we take *N* ≥ 10000, and the maximum number of connections is *K* = 1000 for 10 clusters (Supplementary figure 3d).

The total FF premotor input into an inhibitory neuron i (*i* = 1, …, N) in the motor network is 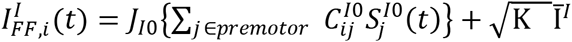. The synaptic strength, *J*_*α*0_, is parameterized as the recurrent synapses: 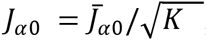, with 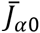 of order unity and 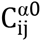 is a random adjacency matrix: 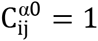 with probability K /N and zero otherwise. Finally, similar to the recurrent interactions, the dynamics of the synaptic variables 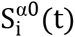 yields: 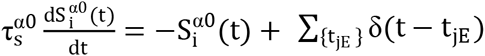 witht_jE_ as the spike times of neuron (*j*, *E*) in the premotor network. For simplicity we take K_FF_ = K.

#### Temporally structured feed-forward input to the motor network

We model the temporally structured inputs to the motor network (HVC to RA input) by including an additional contribution, 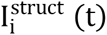, to the FF input received by the neurons in this network. The input, 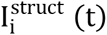, to neuron i, lasts for a duration of 600ms (a typical duration of a zebra finch song motif) repeated 300 times. It consists of a random sequence of On and Off periods, the duration of which are drawn from an exponential distribution with mean 20ms for the On and 70ms for the Off periods. The amplitudes of the input during the On periods are drawn randomly from uniform distribution over an interval [0.1, 0.5]. The input sequences are generated independently for neurons in different functional groups. each group is then divided into 20 (non overlapping) sub-groups such that all the neurons in a subgroup share the sequence. Note that the results depicted in Fig. 4A were obtained by simulating the network *without* structured input.

#### Effectors

The pathway from the motor network to the effectors is topographic. Specifically, we assume that: 1) the number of effectors and the number of groups are equal; 2) a given effector is activated by *M* ≫ *D* neurons in the motor network randomly chosen from one group; 3) Different effectors are activated by different functional groups. We modeled the activation of an effector, E_l_(t)(*l* = 1 … *D*), as a linearly filtered version of the activity of the neurons in the motor network, namely:

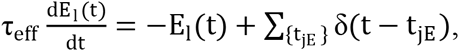

where*τ*_*eff*_ is the effector time constant and the sum is over all spikes emitted by the neurons in the motor network which activates the *l* effector at times t_jE_ < *t*.

#### The vocal organ

We modeled the vocal tract as in Amador et al.^33^. In particular, we did not include the trachea or the Helmholtz filter, as these filters are species-specific and in general will not affect gesture and inter-gesture durations. Two variables activate the vocal organ: tension and pressure. We modeled the pressure variable as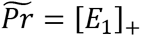, where [*x*]_+_ is a rectified linear function. The tension is modeled as a linear combination of nine effectors: 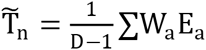, where, W_l_, (*l* = 2, …, *D*), are random weights, W_l_ ~ *N*(0,1). Tension and pressure are then scaled to fit the dynamic range of the oscillating phase (see ref *(33)*):

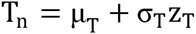

and

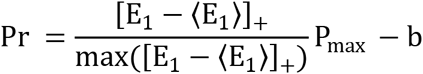

wherez_T_ is the z-score of 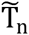 andσ_T_, μ_T_, P_max_ are constant parameters which define the dynamic range and b is a bias which ensures that when there is no pressure the system is at a fixed point. We take: μ_T_ = 0.6; σ_T_ = 0.2; P_max_ = 0.21; b = 0.01. The tension and pressure were then smoothed by a rectangular window of 20ms and interpolated to a sampling frequency of 44,100Hz. We then used the tension and pressure parameters to simulate the Amador et al. model^33^. Finally, to reduce transient effects at the boundaries of the gestures (as a result of crossing the bifurcation) generated by the vocal tract model, the sound signal was taken as the product of the output model and the Pr signal

#### Model parameters

Unless specified otherwise, the parameters used in the simulations were:N = 10,000; K = 400; D = 10; τ_m_ = 10ms; τ_eff_ = 10ms. In the simulations depicted in Fig. 1 synaptic strengths and external FF inputs were: 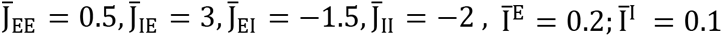 for the premotor as well as for the motor network. All synaptic time constants were 3ms and 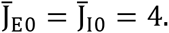 In fig. 3 the parameters in the premotor network were: 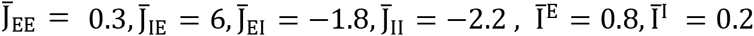 and for the motor network: 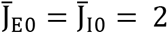 and 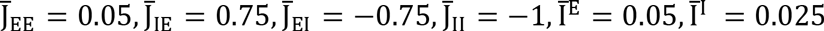. τ_eff_ = 5ms. All synaptic time constants were 3ms except for the premotor-to-motor pathway to the excitatory neurons in the motor network, which represents the slow NMDA synapses in the LMAN-RA pathways (see figure). The parameters used in the simulations depicted in Fig.4,6 were chosen such that the mean firing rates of the neurons in the premotor and motor networks were in agreement with previous experimental data as well as our own data in LMAN and RA. Given these parameters, the average firing rates of excitatory and inhibitory neurons in the premotor network were 14.7Hz and 46Hz, consistent with our data and with previous reports for adult and juvenile finches^10^. The mean firing rates in the motor network are 40Hz for the E cells and 100 Hz for the I cells, as reported for RA neurons^27^. The synaptic time constants of AMPA and GABA_A_ mediated synapses are all taken to be 3ms. NMDA mediated synapses in the premotor-to-motor pathway are modeled in a minimal manner, neglecting their voltage dependence, with very fast (instantaneous) rise and slow exponential decay with time constants of ~100 ms, in line with experiments^30^. We would like to stress here once more that the qualitative behavior of the model is highly robust to changes in all its parameters (see Supplementary Information).

### Numerical simulations and analysis of the results

#### Numerical integration

The dynamics of the model circuit were numerically integrated using the Euler method supplemented with an interpolation estimate of the spike times^53^. In all simulations the integration time step was 0.1*ms*. We verified the validity of the results by performing complementary simulations with smaller time steps.

#### Auto and Cross-Covariance of spike activities

Neuronal spike trains were filtered with an exponential kernel (time constant = 5ms). Auto (AC) and crosscorrelations (CC) of neuronal activities were estimated from the resulting smoothed signals. Population averaged AC and CC were computed over all neurons in the corresponding population. The Pearson CC was defined as the crosscovariance normalized by the autocovariance at zero lag.

#### Measure of synchrony and variability of the effectors

We quantified the degree of synchrony in the activities of the premotor or motor network using the synchrony measure, *χ*(*M*), defined by^54-55^:

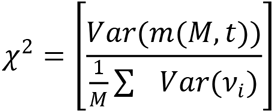

where the sum is over a population of *M* neurons in the network, *v*_*i*_ (*t*) is the instantaneous firing rate of neuron *i* and 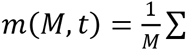 *v*_*i*_(*t*)is the instantaneous firing rate averaged over the population of M neurons. Here, *Var*(*x*(*t*)) denotes the variance of the temporal fluctuations of *x*(*t*) and [*x*] denotes the average over a large number of realizations of the population *S*. For 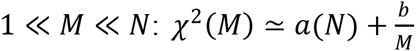, where *a* and *b* are numbers which depend on the network parameters. By definition, the network is in an asynchronous state if *a* vanishes for sufficiently large *N*. In that case pair-wise correlations are small, of the order of 1/N and the population average firing rate is constant in time. In contrast, if *a* converges to a non-zero value for large *N*, the network is in a synchronous state. To quantify the variability of the inputs to the effectors (receiving inputs from M neurons in the motor network), *E*_*l*_(*M, t*) (*l* = 1, …, *D*), we computed the coefficient of variation, *CV*_*eff*_ such that:

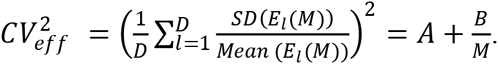

If the motor network is in an asynchronous state, 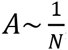 since *E*_*l*_(*t*) is linearly related to the population averaged activity in a functional group *l*.

## Supplementary Information

### Temporal irregularity in strongly connected E-I networks

Asynchronous strong temporal irregularity and spatial heterogeneity in spiking activity of single neurons emerge naturally from the dynamics of recurrently connected neural circuits, in which strong excitation is balanced by strong inhibition (Fig.1b. Supplementary Figure 1a-b; *1-3*). In this regime, neurons exhibit irregular firing as well as large trial-to-trial variability even in the absence of *any* source of noise external to the network. The essence of this temporal irregularity arises from the fact that each neuron receives a large number of “strong” (*1-3*) excitatory and inhibitory presynaptic inputs that cancel each other to the leading order, but the set of inputs is different for each neuron. The latter heterogeneity in the network connectivity results in temporal irregularity of the neuronal activity.

The temporal time scale on which fluctuations in the inputs or outputs of the neurons decorrelate depends on the synaptic dynamics. The theory predicts that for fast synapses this decorrelation happens rapidly (Supplementary Figure 7; (*1-2)*). By contrast, if synaptic interactions are slow (compared to the neuronal integration time) the neuronal activity exhibits slow chaotic rate fluctuations (*4-5)* (Supplementary Figure 7). In both cases, the distribution of the temporal average firing rates of the neurons is well approximated by a log-normal distribution *(6)* (Supplementary Figure 1c). The theory- derived in a completely random networks- also predicts that the averaged pair-wise spatial correlations of the neuronal activity are very weak, on the order of 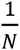 where N is the number of neurons *(3,7-8)* (Supplementary Figure 1d-e). Moreover, in this regime single neuron activity is highly sensitive to temporal fluctuations, as well as to heterogeneities in the recurrent or feedforward connectivity or in small variations in external inputs. These features endow the network with remarkable computational capabilities; e.g., the emergence of selective neuronal responses to stimuli *(9).* These conclusions, combined with experimental results(*3,10-15*), support the idea that the *balanced regime* is a fundamental mode of operation of local cortical networks.

Our paper addresses a fundamental question; namely to what extent the intrinsic temporal variability in recurrent networks can be exploited to drive motor variability. In a more general mechanistic perspective, we investigate how single neuron variability can result in uncertainty in behavior (e.g. in making a decision *(16)*). The fact that in the balanced regime neurons are only weakly correlated should apparently prevent the transfer of internally generated variability at the neuronal level to any downstream system which sums the activity of many fluctuating neurons. In this work we demonstrate that this is not the case. Specifically, we show that spatial correlations can emerge if the circuit generating and transferring the variability to the effectors has a topographic organization. As a result, the circuit can eventually produce robust exploratory behavior. In fact, we show that this mechanism does not require *any fine-tuning of parameters.* In particular, it is robust to the number of neurons N, the number of connections K, as well as to the connectivity in the topographic pathway to the effectors (and the number of neurons projecting to an effector M).

To formulate in a minimal mathematical description the concept of the balanced regime, we consider an unstructured network model of one excitatory (E) and one inhibitory (I) binary neurons receiving strong feedforward inputs from an external population of excitatory neurons *(1-2).* Following van Vreeswijk and Sompolinsky*(1-2)*, the total synaptic inputs into the excitatory and inhibitory neurons can be written:

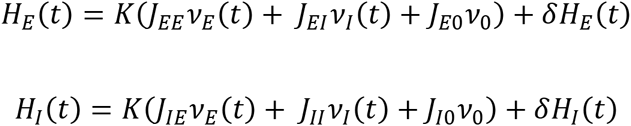

where we denote by *δH*_*α*_(*t*), *α* ∈ {*E, I*} subleading (in large K) contributions to these inputs, which in particular include the temporal fluctuations. Note that the sign of the synaptic strength of the inhibition is negative (*J*_*αI*_ < 0). The variance of these fluctuations is:

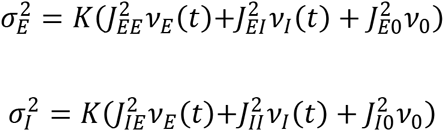

Here, *v*_*E*_(*t*), *v*_*I*_(*t*) and *v*_0_ are the population-averaged firing rates of the E and I neurons and of the external population; *J*_αβ_, α, β ∈ {E, I}and *J*_α0_ are the strengths of the recurrent and feedforward synapses (considered for simplicity to be homogeneous for each synapse type). We also assume that the average number of synapses per neuron, K, is the same for the two populations and for both the recurrent and feedforward synapses and that it is large (the number of synapses is typically 100 or more). Scaling the strengths of the recurrent and feedforward synapses as:

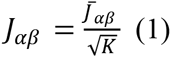

the temporal fluctuations remain finite when K becomes large. Requiring that the mean inputs also remain finite implies:

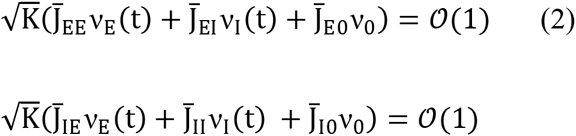

and therefore, for large number of synapses, K, 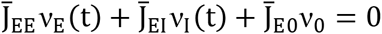 and 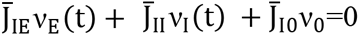. These two linear equations express the fact that, in the two populations, the net inputs into the neurons are comparable to their firing threshold in spite of the fact that taken separately, excitation and the inhibition are much larger than the threshold. These two linear equations uniquely determine the population average firing rates *v*_*E*_ and *v*_*I*_ of the excitatory and the inhibitory populations. The requirement that *v*_*E*_, *v*_*I*_ are positive, yields a set of inequalities which determines the domain of the parameters (synaptic strengths and external inputs) in which the balanced state exists *(1-2)*. Thus, recurrent networks of E and I neurons can operate in the balanced regime for a wide range of parameters without *any fine-tuning* of these parameters. Importantly, because of the scaling of Eq.(1), the amplitude of the temporal fluctuations in the inputs are comparable to the threshold even if K is large. This is why in the balanced regime neurons fire very irregularly. Similar arguments hold for rate-based units *(4)*, as well as for more realistic neuronal models, such as integrate-and-fire or conductance-based neurons *(9)*.

It is important to note that although the average rates of the E and I populations are a linear function of their inputs (see Eq.(2)), the system is highly non-linear and the output of each neuron, as well as the dynamics of sub-populations of neurons (as we will show below), are a non-linear function of their inputs.

### Common feed forward input to all neurons in the motor network is not sufficient to drive motor variability

Transfer of temporal variability from the premotor network downstream to the effectors requires that the motor network operates in the balanced regime. Is it also sufficient? To address this question we investigated the case, where all neurons in the motor network share common inputs from the same group of premotor cells (Supplementary Figure 2a). One might expect that this common feedforward component would tend to synchronize the activity of all the neurons in the motor network. This is not the case. As depicted in Supplementary Figure 2c, the strongly irregular activity of the neurons in the motor network only exhibits very small spatial correlations (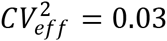). In fact, the strength of the spatial correlations vanishes in inverse proportion to the average number of recurrent connections in the motor network, K (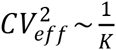; Supplementary Figure 2b-d). This feature is a hallmark of unstructured and strongly connected recurrent networks, where the recurrent inhibition in the motor network *suppresses* the correlations due to shared inputs*(3,7)*. This can be understood heuristically as follows. As long as the matrix *Jαβ*, which characterizes the strength of the interactions in the motor network, is regular (see Eq.(2)) it fully determines the population average firing rates, *v*_*E*_ and *v*_*I*_, as a function of the population average activity of the premotor network (*v*_0_). Therefore, since the latter is constant, *v*_*E*_ and *v*_*I*_ must be constant up to small fluctuations. Hence, the motor network cannot exhibit synchronous activity and neuronal variability cannot be transferred to the effectors.

### Motor variability emerges if the premotor-to-motor projections are topographically organized

If the premotor-to-motor projections are topographically organized, the order 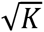 time averaged feedforward inputs to the neurons in the motor network are the same for all functional groups in the motor network. However, their temporal fluctuations, which are of *O*(1), vary from one group to the other. We found that in this case irregular activity with robust spatial correlations emerges in the motor network.

To intuitively understand why this is the case, consider a circuit with two functional groups (Supplementary Figure 5). Denoting by *v*_*E,l*_(t), the time-dependent population average activity of the E population in group l= 1,2, the balanced equations (Eq.(2)) yield two sets of equations for the three populations (one I and two E populations):

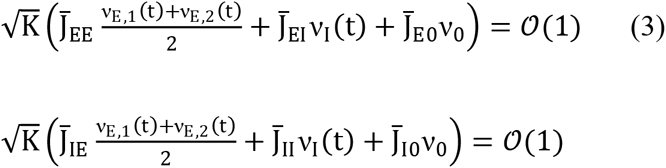

Here we assumed, as in the spiking model we simulated, that the recurrent connectivity is statistically homogeneous over the entire motor network and that the groups are defined solely by the topographic organization of the premotor FF inputs.

The balanced equations imply that 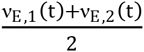 and v_I_(t), the average of the instantaneous population activities of the two groups and the inhibitory population, are constant in time. Therefore, v_E,1_ (t) and v_E,2_ (t) can vary in time, without breaking the balance of excitation and inhibition, provided that they vary in a push-pull manner and are thus negatively correlated.

Numerical simulations indicate that this indeed occurs (analytical proofs will be presented elsewhere). The network recurrent dynamics self-organize such that v_E,1_ (t) and v_E,2_(t) both exhibit significant temporal variations which are driven by the *O*(1) shared fluctuations present in the feedforward inputs. Importantly, the temporal fluctuations in the population activities, v_E,1_ (t)and v_E,2_(t), are large and negatively correlated, in spite of the fact that the shared FF inputs are only weakly correlated between the two groups. This result in an average 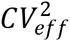 which is large (Supplementary Figure 5b bottom 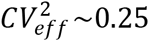) and which is finite (of order unity) even for large N, K and M (Supplementary Figure d-f).

Self-organization of the dynamics also occurs when the number of groups, D, is larger than 2 (Fig.2, Supplementary Figure 3-4). This gives rise to a spatiotemporal pattern of activity in which the firing of the neurons is positively correlated within each of the groups, while the instantaneous firing rate averaged over all the excitatory neurons in the network is constant in time (the E and I population-averaged instantaneous firing rates are constant in time, up to small fluctuations due to finite size effects in K and N).

### The timescales of the fluctuations in the motor network

Our behavioral data shows that vocal babbling temporally decorrelates on a timescale of several tens to a few hundred milliseconds, depending on the species (Fig.7). Moreover, our electrophysiological recordings in singing finches demonstrate that the activity of neurons in RA fluctuates slowly and decorrelates on similar timescales (Fig.5g-i, Fig.6b-c). These timescales are significantly slower than the typical single neuron integration time, raising the question of the origin of such slow timescales.

One possibility is that the recurrent synapses in the motor network are sufficiently strong and their dynamics are sufficiently slow to generate slow chaotic rate fluctuations *(4)* (Supplementary Figure 7b, black curve). However, these fluctuations would be very weakly spatially synchronized (Supplementary Figure 7c, black curve). Thus, spatiotemporal correlations on such slow timescales between neurons projecting to the same effector would be weak and therefore unable to induce highly variable motor behavior (Supplementary Figure 7d, black curve).

Another possibility is that the fluctuations in the shared component of the *feedforward* premotor-to-motor input are slow, resulting in spatially *and* temporally correlated fluctuations at the level of population in the motor network (Supplementary Figure 7c- red and blue curves and Fig.6). This occurs naturally if the premotor-to-motor pathway involves a sufficiently large fraction of slow synapses, e.g. synapses mediated by NMDA receptors, that would low-pass filter the fluctuations generated in the premotor network. This mechanism is depicted in Fig.6 and Fig.7A.

The recurrent dynamics in the premotor network can also contribute to the slowness of the fluctuations in the shared FF inputs to the motor network and therefore to the slow decorrelation of the motor behavior. If the recurrent synapses in the premotor network (the synapses in the E-I-E loop or of the mutual-inhibition) are strong and slow, the dynamics of the premotor neurons will be chaotic (Supplementary Figure 7a- red and blue curves) and their activity will decorrelate on a timescale on the order of the typical (and slow) synaptic time constant*(4)*.

### Correlations between neurons in electrophysiological recordings

In line with our model, the data reported in Figure 5, exhibited positive as well as negative cross-correlations (CCs). However, the majority of the CCs were positive. One should note that our experimental technique for recording single unit activity in singing finches is probably biased toward pairs of neurons located at rather short distances compared to RA diameter. Unfortunately, we cannot accurately report the distance between recorded neurons for two reasons. First, the electrodes (tungsten microwires) are slightly flexible and the distance between their tips (around 100 µm before being moved in the brain) can vary considerably as they are advanced down into RA. Second, post-hoc histological examination was carried out at the end of the experiment, after the electrodes had been advanced several times (with 4 electrode each time), making it impossible to associate a given electrode tract with a given recording. As RA is topographically organized (17-18), it is very likely that our bias to record from nearby neurons resulted in having many pairs recorded from the same functional group.

Our theory can be further validated by showing that pairs of neurons which are far apart, and thus belong to different functional groups, are in general more negatively correlated than more proximal neurons, which are presumably in the same group. As explained above, our single unit data cannot be used for such a validation. Instead, we recorded LFPs in RA of singing zebra finches with fixed implanted electrode micro-arrays (similar to Utah arrays) that had 7 recording sites arranged (with ground and reference electrodes) in a 3x3 lattice with 100 µm minimal distance between electrode tips. As electrodes cannot be moved after implantation, recording single-unit activity from a very dense nucleus such as RA is unfortunately impossible using this technique. However, we were able to carry out noise correlation analysis on LFP signals as we did for single or multi-unit activity. As depicted in Supplementary Figure 6b, LFPs recorded from more distant electrodes were negatively correlated, whereas when the electrodes were closer, positive and negative correlations were observed. Therefore, neurons that were recorded by two electrodes 100 µm apart or less are likely to be more positively correlated, as depicted in Fig.5, than neurons that are far apart, in line with model predictions.

### Correlations measured in RA are not an artifact of a misalignment or variation in duration of the syllables and song motifs

Our electrophysiological data show stronger noise correlations in RA than in LMAN. As the length of the song motif varies slightly from rendition to rendition, misalignment of song-related neuronal activity could lead to spurious correlations, especially in RA which neuronal activity is known to be locked to song (see for example Fig.3). To avoid such spurious correlations we carefully time-warped our spiking data from single and multi-unit recordings based on the timing of single syllable events in the motif produced by the birds (see Material and Methods). An example of how such time-warping changed the precision of song-related firing is illustrated in Supplementary Fig. 6c. As is apparent, Supplementary Fig. 6c clearly shows that spikes are more aligned to the song motif following the time warping procedure. To assess the contribution of time jitter from behavioral misalignment to the noise correlations in RA, we re-calculated the noise correlations without any time-warping. Surprisingly, there was very little change in the level of correlation with and without time-warping.

We also estimated how much misalignment and bad time-warping could contribute to spurious correlations by shifting together the spikes of the two simultaneously recorded neurons presented in Fig.5h on each rendition by a time jitter in the range 0-500ms (Supplementary Figure 6d). The correlations did increase with time jitter, as expected, but this only occurred at very large jitters above 150ms. Interestingly, the amount of correlations between the two neurons hardly changed when the jitter was up to 50 ms, far more than the estimated elasticity of the zebra finch song (around 5ms at most for a given syllable, see (19)). Therefore, for typical jitters, misalignment of the motifs did not contribute to the noise correlations measured in RA.

### On alternative mechanisms

Three key hypotheses underlie our work: (1) The variability of the babbling is produced by a circuit in the CNS comprising large networks of strongly coupled neurons firing a highly irregular manner (2) These fluctuations in the activity are transferred to the effectors which sum up inputs from a rather large number of neurons, on the order of 100 or more. (3) The mechanism underlying babbling should be robust, in particular with respect to variations in the connectivity parameters.

It should be noted that if we relax Hypotheses (2); i.e, if we assume that the projections to the effectors only involve a small number of RA neurons (very sparse projections to the effectors) topographic projections from LMAN to RA are no longer necessary for the circuit to generate fluctuations in the inputs to the effectors. However, the amplitude of the latter depends crucially on the connectivity which means relaxing Hypotheses (3).

In fact, the last two hypotheses imply that fluctuations in the activity of the neurons should become correlated at some stage in the circuit before their transfer to the effectors and that this should be a *collective* phenomenon.

In all four species we considered in our behavioral study the ACE of the babbling signal in juveniles lacked an oscillatory component, and the gesture duration distribution was well approximated by an exponential. This means that the process of generating gestures during babbling is very broad band, and has no substantial oscillatory components. The electrophysiological data suggest that the fluctuations in neural activity in RA were also very broad band without significant oscillatory components (see the autocorrelations and cross-correlations depicted in Figures 5-6). We thus need to look for network mechanisms that can account for correlated neuronal activity exhibiting strong fluctuations without having specific frequencies in the power spectrum. This is a non-trivial constraint. In fact, in previously investigated mechanisms for robust correlated activity, the latter stems from the emergence of oscillatory collective modes in the network dynamics (for review see (20)). This is the case in the mechanisms for spike-to-spike synchrony, as well as well as in those in which synchrony emerges from firing-rate instability (see e.g. (21)). In most of these models, the oscillation phase basically remains constant over many cycles, unlike what is observed experimentally. Solutions to this problem have been proposed. They all rely on synchronous chaos and lead to temporal irregular fluctuations in the population activity (22-25). However, the spectrum of these fluctuations- although broad - is also peaked in some frequency ranges (the gamma range in the papers cited above). This is because synchronous chaos emerges from a destabilization of synchronous patterns of activity which oscillate at a frequency in this range. To the best of our knowledge, our mechanism is the first to exhibit irregular synchronous activity which to a large extent is robust to changes in the number of neurons and in the average number per neuron of feedforward and recurrent connections.

We also argue that NMDA receptors (NMDAR) in the projections from LMAN to RA can account for the rather slow time constant over which babbling decorrelates. However it is also possible that low-pass filtering occurring in RA or downstream to it due to slow neuronal dynamics (such as adaptation currents) or slow muscle responses could also contribute. Such low-pass filtering would lead to a slower behavioral output than would be predicted from the NMDAR time constant alone. However, in the case of zebra finches, the NMDAR kinetics measures in LMAN-RA synaptic inputs in juvenile zebra finches (26) fit satisfactorily with the timescale of babbling vocalization we report, suggesting minimal low-pass filtering downstream. Moreover, muscle dynamics has been reported to be fast in the bird syrinx (<5ms, see 27-28), and even if young birds have slower muscles it is hard to imagine that their dynamics would involve timescales as long as 50-100ms. This point however needs to be confirmed experimentally.

#### Note regarding Supplementary Figure 8

To calculate the transformation *C*_*g*_ (Δ) = *F*_*g*_ (*C*_*ξ*_ (Δ)) given in Supplementary Figure 8E we assumed a Gaussian Process *ξ(t)*, with *lim*_*t*_*→*_∞_〈*ξ*(t)〉 = 0 *lim*_*t*_*→*∞〈*ξ*(t)*ξ*(t + Δ)〉 = *C*_*ξ*_(Δ) and *c*_*ξ*_*(0)* = 1. Consider the threshold-power-law function: *g*_ϵγ_(*x*) = *g*(*x* − ϵ)Θ(*x* − ϵ)with *g*(*x*)=x^γ^. The AC of the process {*g*_ϵ*γ*_(*ξ*)} is then: 〈*g*_ϵγ_{ξ(t))*g*_ϵγ_{*ξ*(t + *τ*))〉 — 〈*g*_ϵγ_{*ξ*(*t*)〉^2^ = *c*_*g*_(*τ*). The first and second moments are:

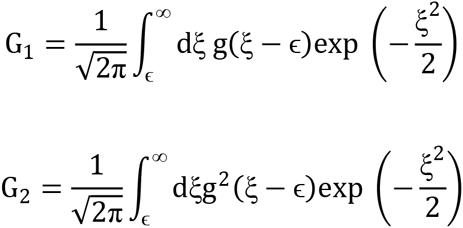

and the correlation function:

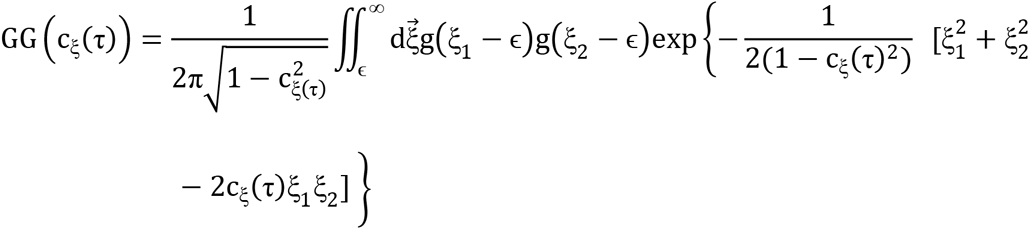

We then numerically calculate the 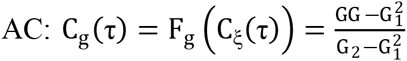. Note that for *ϵ* = ∞ and *g*(*x*) = *x* we get *C*_*g*__*ϵ*(*τ*) = *C*_*ξ*_(*τ*), which is the identity function of *F*_*g*_, plotted in dashed line in Supplementary Figure 8E.

## Supplementary Figures

**Supplementary Figure 1.**
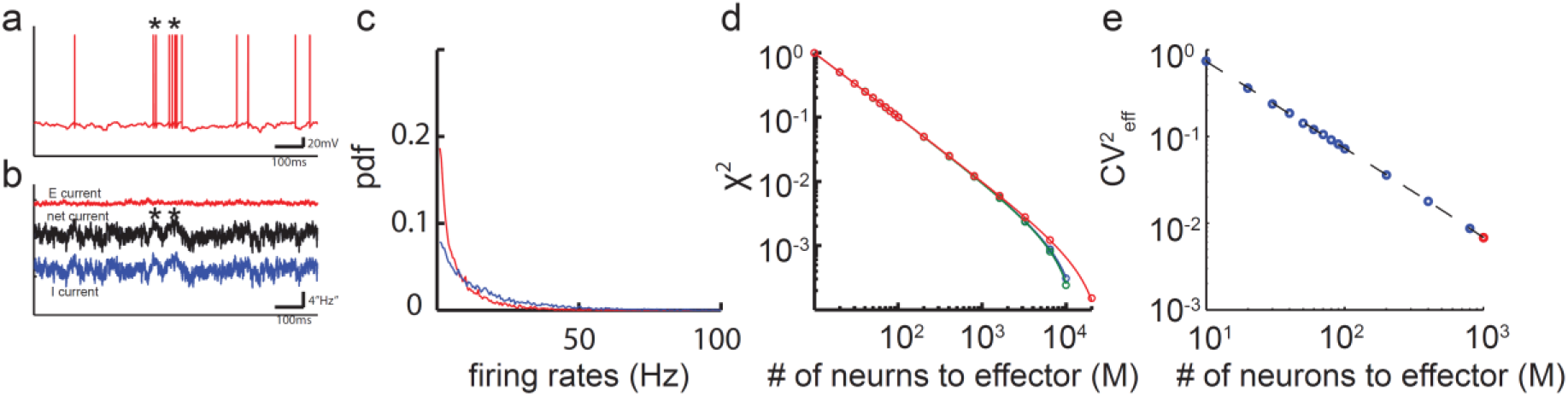
**In the balanced excitation-inhibition regime spiking is irregular, asynchronous and time-averaged firing rates are heterogeneous.** Network parameters are as in Fig.1 **a.** Voltage trace of one excitatory neuron in the motor network. The temporal irregularity in the action potentials is intrinsically generated by the recurrent network dynamics since there is no external noise. **b.** Total excitatory (red), inhibitory (blue) and net (E+I, black) inputs to the same neuron as in(a).The excitation and the inhibition taken separately are large relative to the threshold, but the mean and the fluctuations of the net input (black) are comparable with the threshold. Suprathreshold fluctuations in the net inputs induce irregular spikes (stars in a,b). **c.** The single neuron firing rates in the excitatory (red) and inhibitory (blue) populations are highly heterogeneous. Their distributions are long-tailed. Note that this heterogeneity stems solely from the recurrent dynamics of the network since in each population all the neurons are identical in our model. **d.** Measure of synchrony, *χ*^2^(*M*), as a function of population size, M, in log-log scale (see Materials and Methods). Blue: N=10000,K=400. Green: N=10000, K=800; Red: N=20000, K=400. Blue and green lines are almost indistinguishable. In all cases *χ*^2^~*b*/*M*, up to deviations for *M* ≈ *N*. **e.** 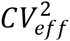 decreases as 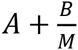 with A≈0 (dashed line). Circles: simulations. Red dot corresponds to the parameters used in Fig1.

**Supplementary Figure 2.**
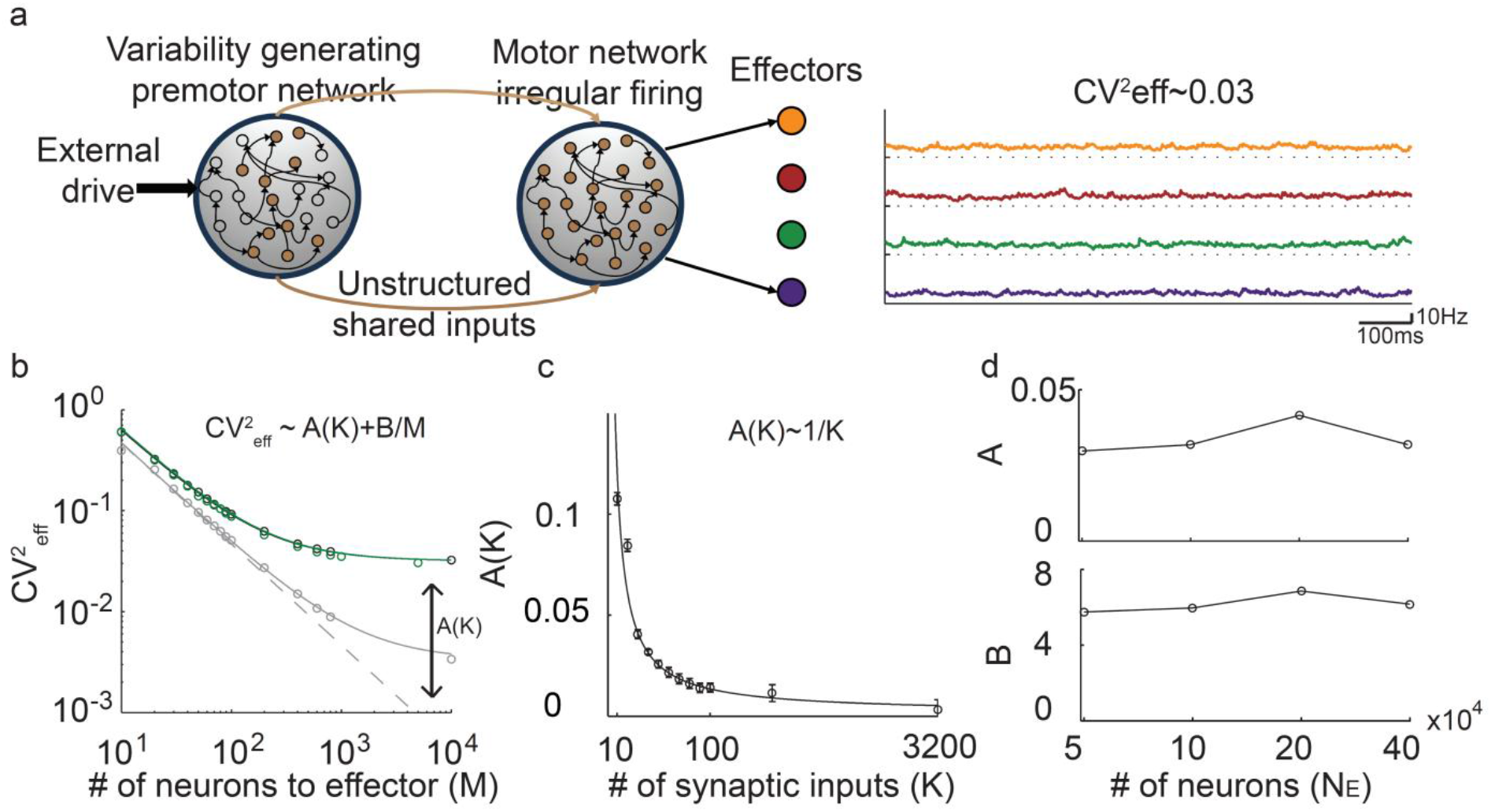
**The activity in the motor network is weakly correlated when all neurons in the motor network share the same premotor inputs. a.** Left: a subset of neurons in the premotor network projects to all neurons in the motor network, thereby all neurons in the motor network share the same feedforward input. Right: the fluctuations in the inputs to the effectors are higher than in Fig.1b, but still very small 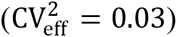 and their magnitude vanishes when the number of synaptic inputs increases. **b-d.** 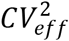 is well fit to 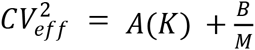for large *K*. In (b): Green: *N*_*E*_ = 40000, *K* = 400; Black (almost coincide with the green): *N*_*E*_ = 10000, *K* = 400; Gray: *N*_*E*_ = 40000, *K* = 3200. In (d) K=400. A and B barely depend on *N*_*E*_. From (b-d) it can be concluded that*A*(*K*) ~ 1/*K*, namely the synchrony in the network is very weak.

**Supplementary Figure 3.**
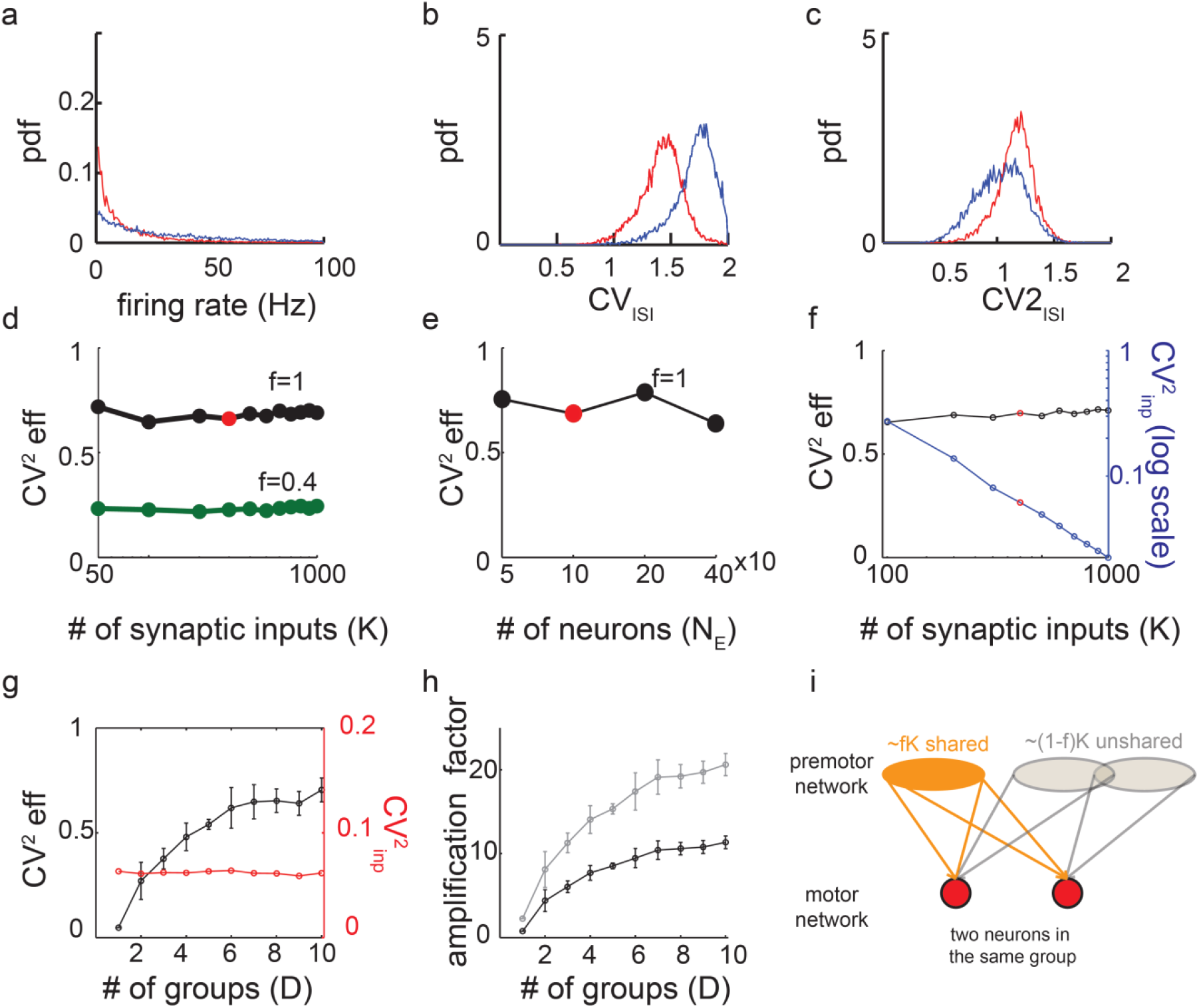
**When the circuit is topographically organized the activity in the motor network is irregular, heterogeneous and the inputs to the effectors exhibit robust temporal fluctuations.** Parameters are as in s. **a.** Distribution of firing rates of E and I neurons in the motor network. **b.** Distributions of *CV*_*ISI*_ for the E and I neurons. Neurons fire irregularly with high *CV*_*ISI*_. **c.** Distributions of 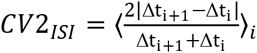 for the E and I neurons in the motor network. Here, Δt_i_ is the i’th ISI and the average is over all the ISIs. CV2 measures the local variability in non-stationary spike trains. Neurons are firing irregularly with CV2 ~ 1. **d-e.** The coefficient of variation of the inputs to the effectors depends only weakly on (d) the number of connections (feedforward and recurrent) and (e) the number of neurons in the network. Red dot corresponds to the parameters used in Fig2.a-e. **f.** The CV of the inputs to the neurons (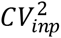, blue) and the inputs to the effectors (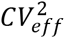, black) plotted vs K. 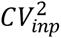 decreases with K while 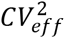 remains essentially constant. This results in an amplification of CVs (see Fig.2g). Red dot: K=400 as in Fig2.a-e. **g.** 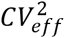 (black) and 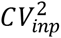 (red) vs. number of functional groups. The average correlations in neuronal activity within a group increases with the number of groups. This stems from the fact that increasing the number of groups results in effectively narrowing the spatial extent of the correlations in the FF inputs with respect to the footprint of the recurrent connectivity in the motor network (see also Fg.2h-i).**h.** Amplification factor 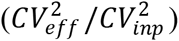 increases with the number of functional groups. Black: *τ*_*eff*_ = 10*ms* as in Fig. 1-2. Gray: *τ*_*eff*_ = 3*ms*.**i.** Cartoon of the FF projections from the premotor network to the motor network. The two neurons in the motor network are in the same functional group. By construction, the two neurons receive *fK* inputs from the same neurons in the premotor network (orange, shared inputs). The neurons also receive inputs drawn randomly and independently from excitatory neurons in the premotor network with probability(1 − *f*)*K*/*N*. (grey, ‘unshared’). The probability that the two neurons also have common inputs in this set is on the order of *K*^2^/*N*^2^ which is small as *N* ≫ *K*. The overlap between the ‘unshared’ and ‘shared’ sets of inputs is not represented.

**Supplementary Figure 4.**
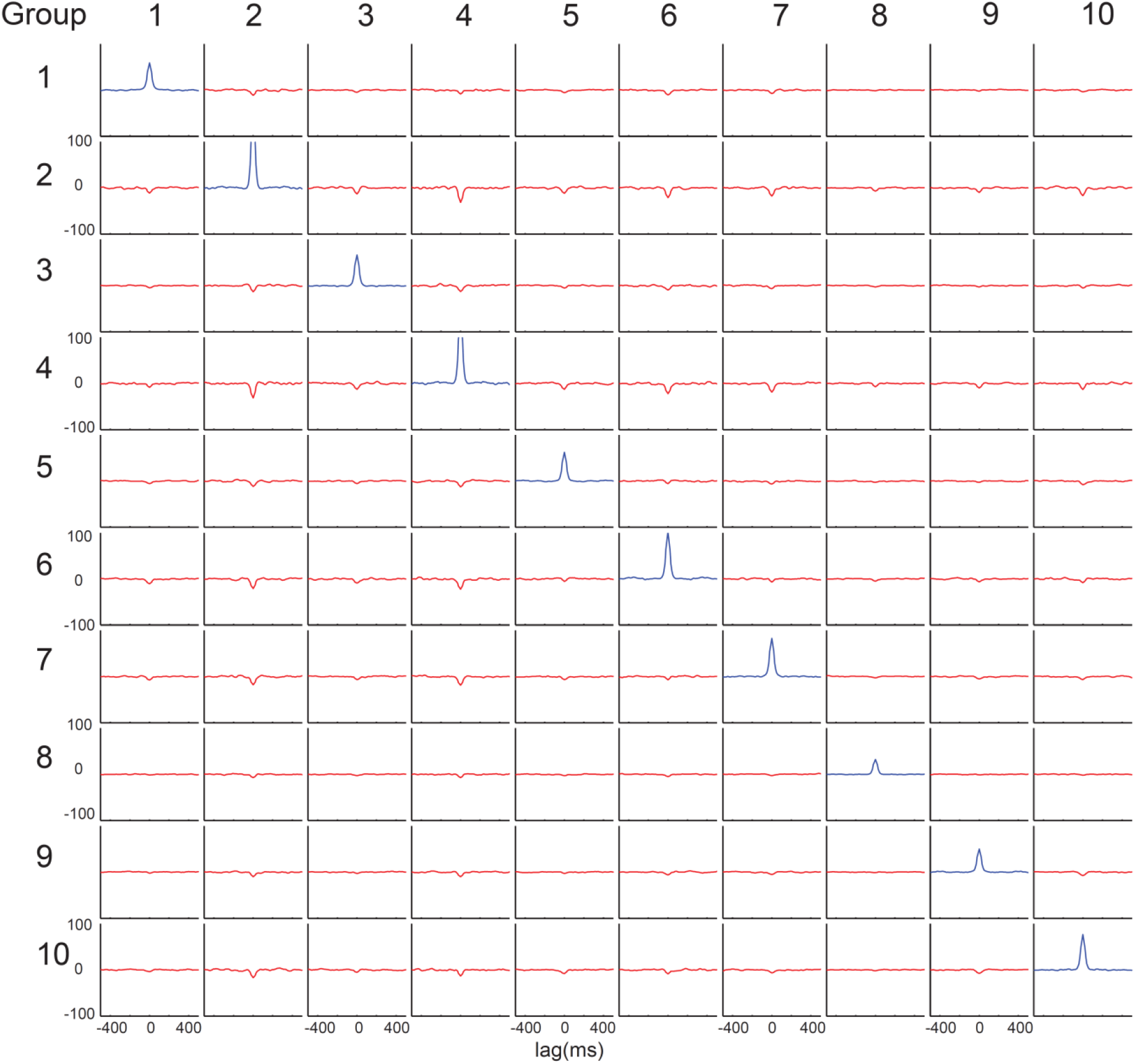
**Crosscorrelations of the activity within and between functional groups in the motor network.** The population average activity of each functional group was smoothed with an exponential sliding window of 10ms. The autocorrelations (diagonal, blue; not normalized) and the crosscorrelations (off-diagonal, red; not normalized) are plotted. Note that the crosscorrelations within a group are positive and much stronger than correlations across groups, which are in general weak and negative.

**Supplementary Figure 5.**
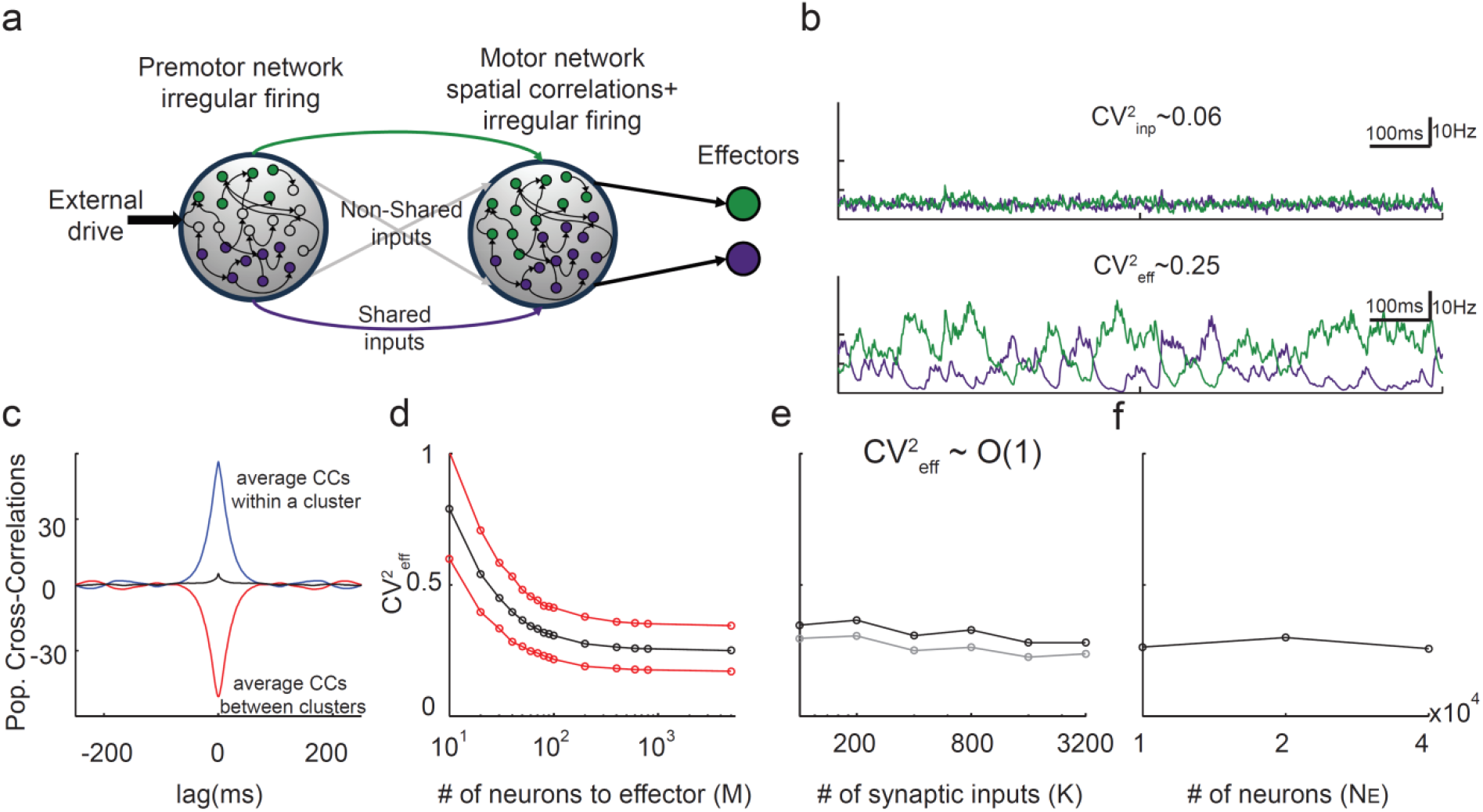
**Push-pull dynamics and spatial correlations for two functional groups.a.** Architecture of the circuit. **b.** Top: Feedforward inputs to neurons in each functional group in the motor network (top). Bottom: Input to the effectors. M=1000. Note that 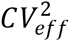 is much larger than 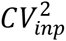. **c.** Population average crosscorrelations for neurons projecting to the same effector are positive (blue), while average correlations between neurons projecting to different effectors are negative (red). Black: population average cross correlations over all E neurons in the motor network. **d-f.**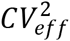 converges to a non-zero value, which only very weakly depends on the number of connections (e) or neurons (f) in the network. **d.** Red: 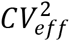 for the two effectors; Black: average 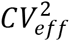 of the two effectors. **e.** Black: M=100; Gray: M=5000. **f.** M=400. Results for M=800 are similar and are not shown.

**Supplementary Figure 6.**
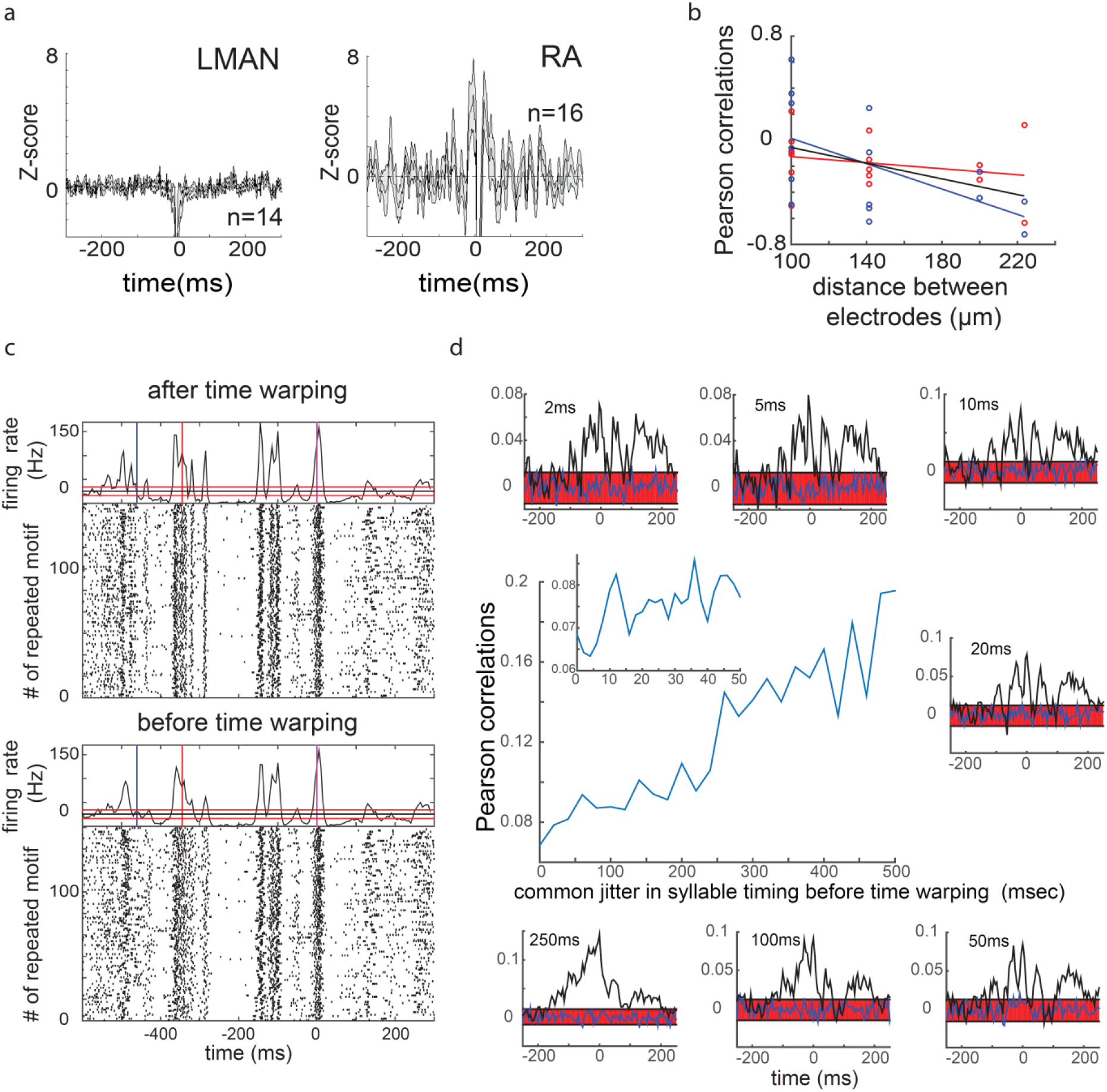
**Additional electrophysiological support for the measured correlations in LMAN and RA. a.** Correlations of LMAN or RA single-units with the multi-units background activity. Left: Spike triggered average (STA) with the background multi-unit envelope in LMAN during singing. The background envelope of multi-unit activity recorded simultaneously with a single-unit on the same electrode is smoothed with a 5-ms Gaussian window. The average motif background activity is subtracted from the singing-related background. STA with this residual signal is then computed for each single-unit recording separately, and the average z-score±s.e.m is shown. Note that a ~10-ms artifact around the spike time is due to the spike subtraction. Right: same as in the left panel, but for RA neurons. **b.** Noise correlations between LFP recordings vs. the distance in the recorded site (see Material and Methods). Note that correlations tend to be negative between sites that are far apart. Each circle denotes a pair of recording sites, and the data were recorded in two birds (red and blue). Solid lines: linear fit for bird 1 (red; slope -0.23; R^2^=-0.056; n.s), for bird 2 (blue; slope -0.56; R^2^ = 0.31; p=0.001) and for the two together (n=15; black; slope -0.43; R^2^=0.18; p=6 10^−14^). **c.** Raster plot of the example RA neuron also plotted in Fig.3a before (bottom) and following (top) time-warping. Note the improvement in the alignment of the spikes to the song motif. **d.** Noise correlations across the two RA neurons depicted in Fig.6b and 5h without time warping and with a common jitter (ranging from 2 to 500ms) applied to the syllable times across renditions of the motif (see Supplementary Information). Main figure: Pearson correlations vs. the amount of jitter. Inset: zoom-in on 0-50msjitters shows that the jitter does not dramatically change the level of correlation. Figures around the main figure: noise correlations as depicted in Fig.5h, but with an increasing time jitter (clockwise).

**Supplementary Figure 7.**
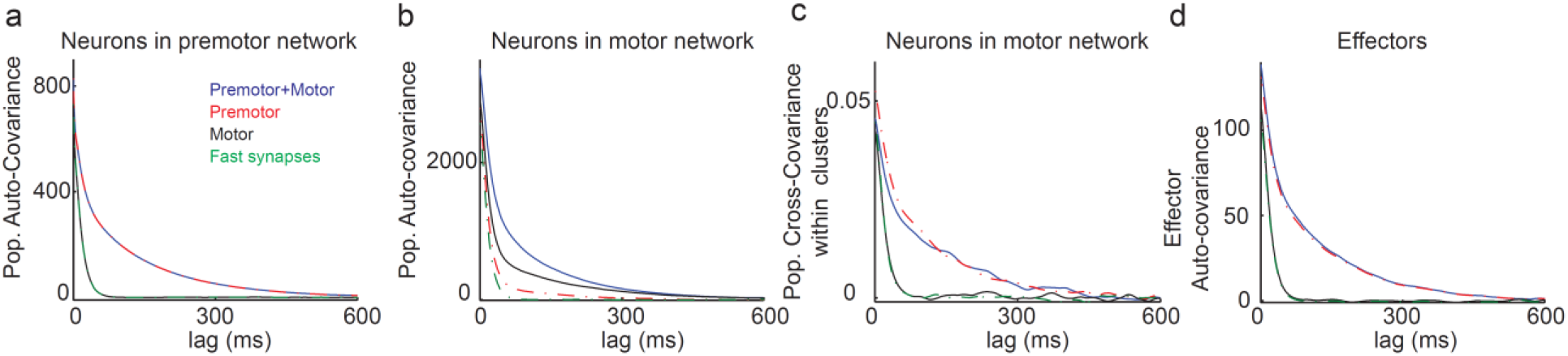
**Various mechanisms generating slow fluctuations in the motor network and in the effectors. a.** Population average autocovariance (not normalized) of neurons in the premotor network. **b.** Population average autocovariance (not normalized) of neurons in the motor network. **c.** Population average crosscovariance (not normalized) for neurons in the motor network in the same functional group. **d.** The autocovariance (not normalized) of the input to the effectors. In all panels the color code is as follows. Green: all synapses are fast (synaptic time constants are 3 ms) and the strength of the mutual inhibition is 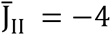. The decorrelation of the fluctuations is fast in the premotor and motor networks (a-b) and in the effectors (d). Red: the mutual inhibition in the premotor network has two components, one fast 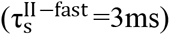 and one slow 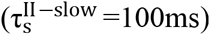. For both components K=400 and their strength is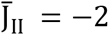. All other synapses are fast. The premotor network now exhibits asynchronous chaotic rate fluctuations which decorrelate on a timescale (a) on the order of the synaptic time constant of the slow inhibition^15^. Both the synchronous activity in the motor network (c) and the input to the effectors (d) decorrelate on the same timescale. Black: the mutual inhibition in the *motor* network has two components, one fast and one slow with the same strength, 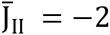. All other synapses are fast. The *motor* network now exhibits slow asynchronous chaotic rate fluctuations on the slow timescale of the inhibition (b) and fast synchronous fluctuations driven by the premotor network (c). Therefore, the fluctuations in the input to the effectors (d) are fast. Blue: the mutual inhibition in both networks has two components, one fast and one slow with the same strength is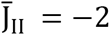. In all panels 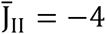 in both networks, unless stated otherwise. Other parameters: 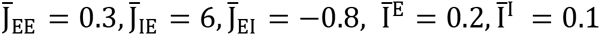 for the premotor network; 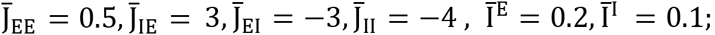 and 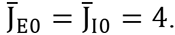.

**Supplementary Figure 8.**
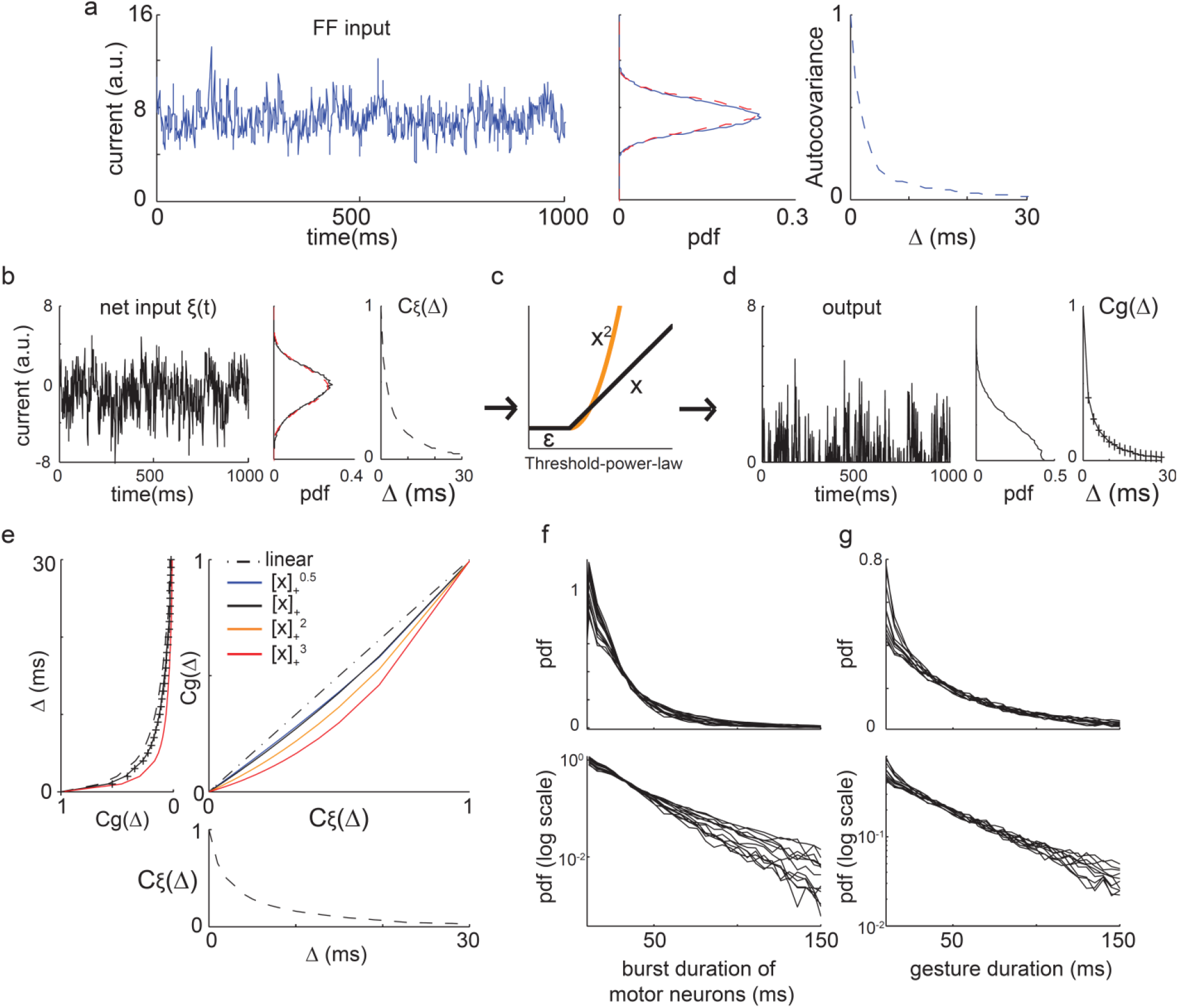
**Statistics of a wideband Gaussian process after a rectified power-law transformations.** In (a), (b) and (d): Left: trace of the process. Middle: marginal distribution of the process. Right: Autocovariance (AC) of the process depicted on the left panel. Parameters in a-b are as in Fig.7a, but with D=2. **a.** FF input to a typical neuron in the motor network. Red: the marginal distribution is well fit by a Gaussian. **b.** Net input to a typical neuron in the motor network. Red: the marginal distribution is well fit by a Gaussian (red). *C*_*ξ*_ (Δ): AC of the process shown in left. **c.** Rectified power-law function 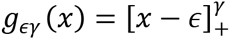. Black: *γ* = 1. Orange: *γ* = 2. **d.** The process in (b) following a rectified linear transformation: *g*_01_(*x*). *C*_*g*_(Δ): AC of the process shown in left. The pdf was estimated after excluding points smaller than 0.1. **e.** Bottom: AC of a Gaussian process {ξ_*t*_} with AC as in (b). Middle: Transformation between *C*_*ξ*_(Δ) to*C*_*g*_(Δ) for different shapes of rectified power-law functions with different exponents. Left: AC of *g*_*ϵγ*_(*ξ*). Dashed black: *C*_*ξ*_(Δ). Black: *C*_*g*_(Δ)for *ϵ* = 0, *γ* = 1. Orange: *C*_*g*_(Δ)for *ϵ* = 0, *γ* = 2. Red: *C*_*g*_(Δ)for *ϵ* = 0, *γ* = 3. Black crosses: AC of the process in (d). **f.** Top: Distribution of burst duration for 14 randomly chosen neurons in the motor network. Bottom: y-axis is in log scale. **g.** Top: distribution of ‘gesture’ durations with a simplified thresholding of the input to the effectors (*γ* = 1). Bottom: y axis is in log-scale. Parameters in f-g are as in Fig.7a and with 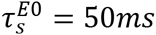.

**Supplementary Figure 9.**
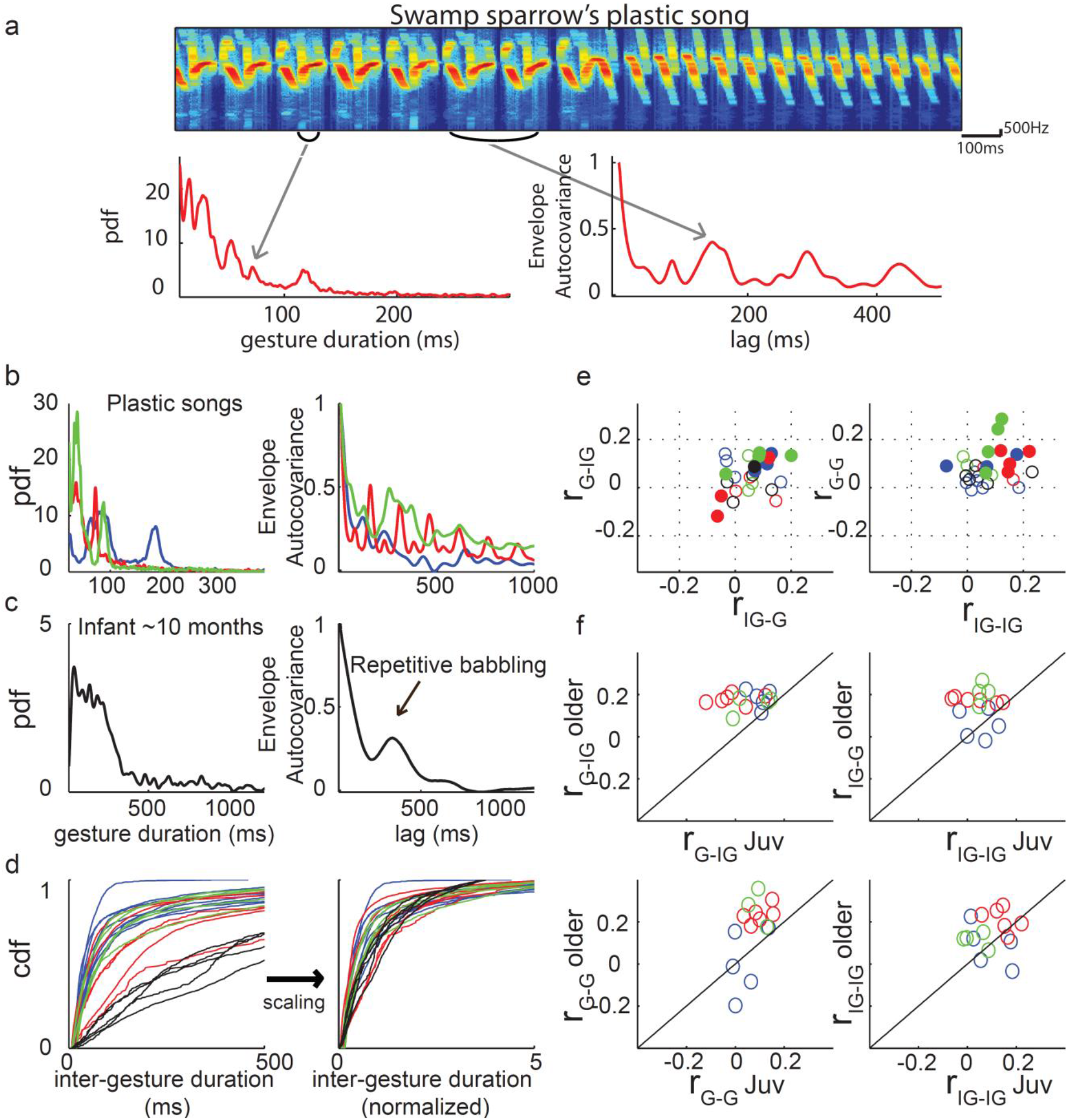
**Statistics for gesture and inter-gesture duration in young and older juveniles.** Color code as in Fig.7. **a**.Temporal structure of Sw plastic song (322-364 DPH). Top: Spectrogram of vocalizations. Bottom: probability density function (pdf) of vocal gesture durations (left) and averaged autocovariance of the envelope (ACE; right). The peaks in the distribution and the ACE express the temporal stereotypy of the song. **b.** Stereotypy of plastic songs and diversity of songs in different species as captured by the gesture duration distributions and ACEs. Examples of gesture distribution (left) and ACE (right) of plastic songs. Note the peaks in the distributions as well as in the ACEs (Sw: 337-379 DPH; Zf: 73 DPH; Ca: 284 DPH, day post hatched). **c.** Gesture duration distribution (left) and ACE (right) of a 10 month old infant, at the beginning of a repetitive babbling period (also called “canonical babbling”, with repetition of the constant-vowels, e.g. ba-ba-ba). **d.** In babbling juveniles, most of the variability between species in the CDF of the inter-gesture duration is accounted by a scaling factor of the time (as is the case for the gesture duration distributions, see Fig7.). **e.** Babbling juveniles. Left: Pearson correlations between the duration of an inter-gesture and the consecutive gesture duration (*r*_*G–IG*_) against the correlation between the duration of a gesture and the consecutive inter-gesture duration (*r*_*IG–G*_). Right: same for consecutive gestures (*r*_*G–G*_) against consecutive inter-gestures (*r*_*IG–IG*_). Full circles: significant non-zero correlations (for both statistics; permutation test; p<0.01). Note that in all cases correlations are close to zero with a slight tendency to be positive, probably due to global tempo changes. **f.** The Pearson correlations increase with age. Correlations for later plastic songs are larger than during babbling for the same individuals.

**Supplementary Figure 10.**
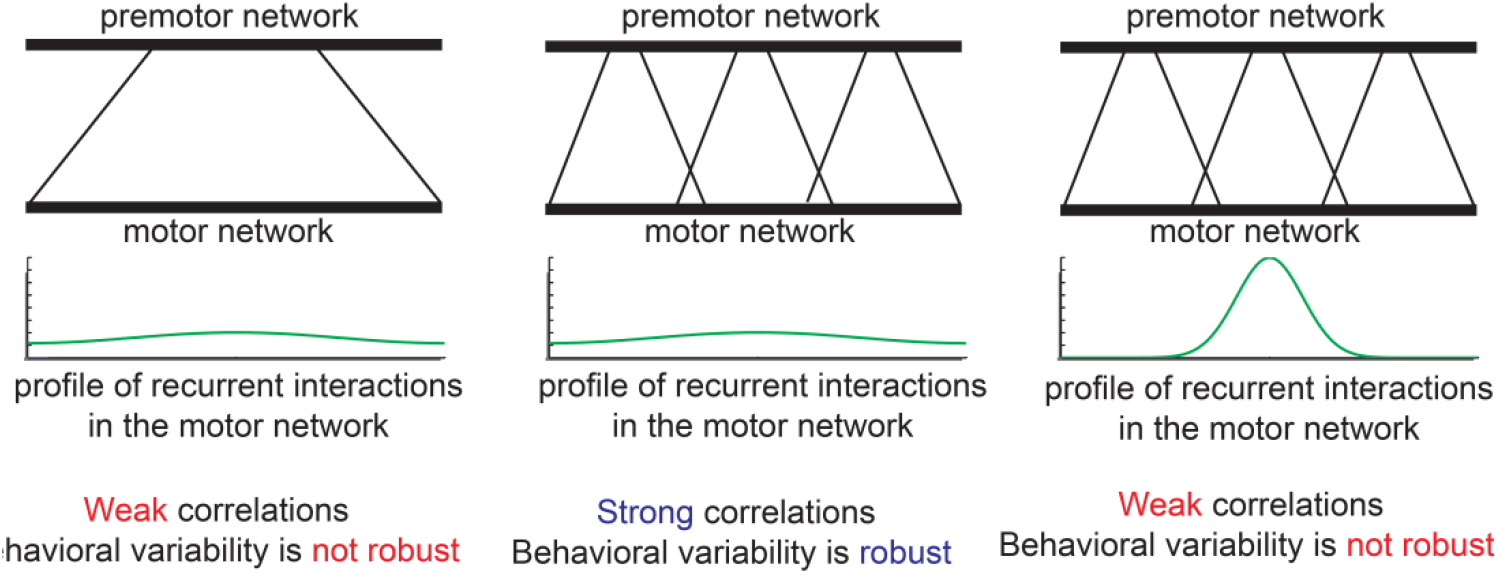
**Robust behavioral variability emerges from the interplay between topographic organization in the premotor-to-motor pathway and the recurrent dynamics in the motor network.** The architecture of the premotor-to-motor pathway (top) and the footprint of the recurrent connections within the motor network (bottom) is depicted in each of the three panels. Neurons in the motor network can develop highly robust correlations when the premotor-to-motor pathway is topographically organized and the recurrent connectivity in the motor network is sufficiently wide.

